# Molecular basis of cooperative assembly of the Ndc80-Ska kinetochore complex on microtubules

**DOI:** 10.64898/2026.04.18.719381

**Authors:** Yiming Niu, Daniel Martsch, Sabrina Ghetti, Jason Mak, Oliver Hofnagel, Daniel Prumbaum, Hironori Funabiki, Andrea Musacchio

**Author notes:** These authors contributed equally (Yiming Niu, Daniel Martsch, Sabrina Ghetti, Jason Mak).

## Abstract

Successful chromosome segregation depends on robust kinetochore-microtubule attachments. The outer kinetochore load-bearers Ndc80 and Ska complexes functionally cooperate, but the molecular basis of their interaction remains elusive. Here, we combine cryo-EM and functional investigations of Ndc80:Ska on microtubules. Ndc80 forms longitudinal arrays along single protofilaments using two modules. The HEC1 N-terminal tail stabilizes interactions between microtubule-binding heads regulated by Aurora B. The HEC1 loop, away from microtubules, organizes Ndc80 coiled-coils into stacks matching the periodicity of tubulin subunits. SkaC binds to a previously unknown interface of Ndc80 as well as to microtubules, simultaneously stapling tubulin dimers longitudinally and neighboring protofilaments laterally. Our work demonstrates how several weak interactions of a small number of individual complexes are harnessed to generate a robust and regulated kinetochore coupler.

## Introduction

Equal segregation of sister chromatids during mitosis relies on bioriented chromosome attachments to spindle microtubules. The kinetochore, a large and tightly controlled macromolecular assembly on chromosomes, harnesses microtubule-binding proteins that can track the plus-end of depolymerizing microtubules as well as resist tension (*1, 2*). This further stabilizes the bioriented microtubule attachment (*3-7*). The 4-subunit Ndc80 complex (Ndc80C), comprising HEC1 (also called NDC80), NUF2, SPC25 and SPC24; and the 3-subunit Ska complex (SkaC), comprising SKA1, SKA2, and SKA3 (**Figure 1**A) are essential for bioriented attachments. Both Ndc80C and SkaC can bind microtubules autonomously in vitro, but neither is sufficient for the formation of stable load-bearing end-on kinetochore-microtubule attachment (*7-15*). In addition to binding microtubules, Ndc80C acts as a direct kinetochore receptor of SkaC (*5, 9, 16, 17*). After recruitment, SkaC is believed to establish a functional unit with Ndc80C to bind microtubules cooperatively and track dynamically polymerizing and depolymerizing microtubule ends (*4, 5, 8, 16-23*). How the SkaC is recruited to Ndc80C, and how its cooperative assembly ensures the extraordinary accuracy of chromosome segregation remain fundamental unresolved question. Here, we report a cryogenic electron microscopy (cryo-EM) analysis of the human full length Ndc80 and Ska complexes on microtubules that sheds new light on these questions.

**Fig. 1.**
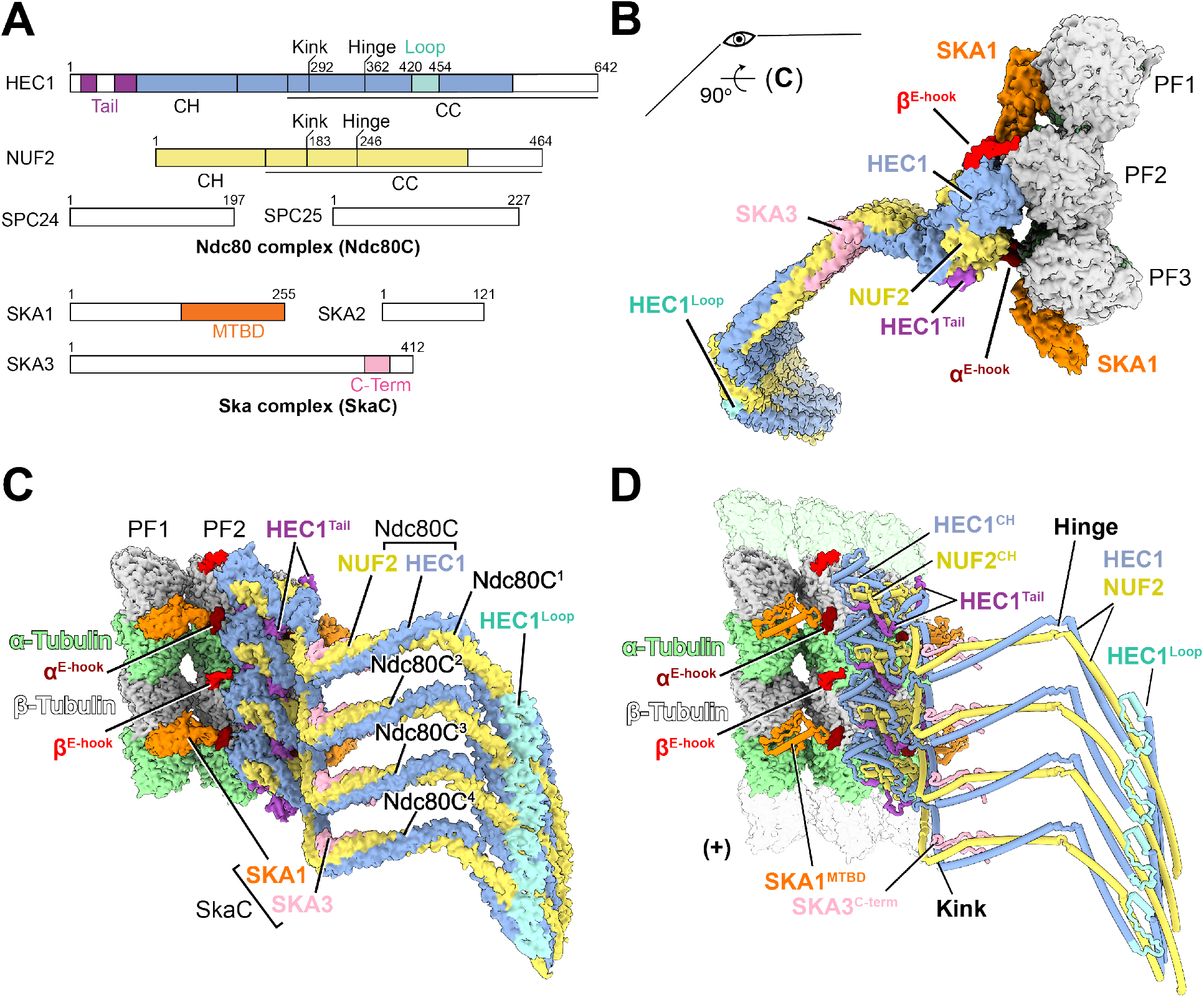
Cryo-EM structure of Ndc80C:SkaC co-assembled on the microtubule. **(A)** Schematic depiction of human Ndc80C and SkaC subunits. Resolved cryo-EM densities are color-coded: HEC1 (blue), HEC1 N-terminal tail (HEC1^Tail^, purple), HEC1 loop (HEC1^Loop^, cyan), NUF2 (yellow), SKA1 microtubule-binding domain (SKA1^MTBD^, orange), SKA3 C-terminal domain (SKA3^C-term^, pink). (**B and C**) Cryo-EM density map of Ndc80C:SkaC on the microtubule. **(B)** Top view from the microtubule minus end. Ndc80C bound to a central protofilament (PF2) via HEC1 calponin homology (CH) domain is flanked by two SKA1 subunits of SkaC on PF1 and PF3. The C-terminal E-hooks of α- and β-tubulin are indicated as α^E-hook^ (red) and β^E-hook^ (brown). **(C)** Side view. Four Ndc80C protomers on both α-tubulin (green) and β-tubulin (grey), are linked by HEC1^Tail^ and HEC1^Loop^. SKA3^C-term^ (pink) associates with the kink of HEC1:NUF2 coiled-coil (CC). SKA1^MTBD^ bound at the inter-α/β-tubulin dimer interface on PF1 also contacts to α^E-hook^ of adjacent PF2, where two neighboring HEC1 CH domains engage. (**D**) Atomic model of (C). The periodicity of tubulin dimers was displayed as transparent α-tubulin (top) and β-tubulin (bottom) for illustrative purposes. Microtubule plus-end (+) is indicated.

### Overall view of the Ndc80:Ska complex

We determined the cryo-EM structure of full-length human Ndc80C and SkaC decorating microtubules. To promote Ndc80C-SkaC interaction, SkaC was prephosphorylated in vitro with active CDK1 kinase (*5, 16, 17, 24*). Extensive 2D and 3D classifications directed on complexes decorating 13-protofilament microtubules, combined with focused masking and local refinement allowed us to obtain two maps at ∼4.0 and ∼3.3 Å resolution (**Figure S1, S2** and Methods). Collectively, these maps allowed unequivocal fitting of models for the segments of Ndc80C and SkaC that are displayed with colors in the schemes in **Figure 1**A, with the rest, displayed in white, being invisible or visualized with lower confidence.

Ndc80C binds to the microtubule lattice *(gray*, α-tubulin; *light green*, β-tubulin) through its head consisting of calponin-homology (CH) domains in HEC1 *(blue)* and NUF2 *(yellow)* (*25, 26*). Individual Ndc80Cs stack onto each other in compact longitudinal arrays (i.e. along the same microtubule protofilament), occupying both α- and β-tubulin (**Figure 1**B-C), as previously observed in a lower-resolution structure with Ndc80C^Bonsai^, an engineered Ndc80C lacking most of the coiled-coil (*26*). Consistent with a recent structure of the HEC1:NUF2 subcomplex (*27*), our structure illustrates the role of the HEC1 N-terminal tail *(purple)* in the stabilization of the stacks.

Unlike previous structures, however, full-length Ndc80C did not exhibit lateral oligomerization on microtubules. A crucial involvement of the HEC1:NUF2 coiled-coil region in the stacking mechanism is now revealed. In an arrangement with an unanticipated kink upstream of the canonical hinge (**Figure 1**D), the HEC1 loop regions (cyan) mirror the longitudinal stacking rule of the microtubule-binding heads, exhibiting a ∼40 Å (1Å = 0.1 nm) period similar to that of tubulin monomers (**Figure 1**C and **Figure S2**C). They form a continuous array that stabilizes a coiled-coil bundle that aligns with the microtubule axis. This orientation predicts that the kinetochore-binding modules of Ndc80C (SPC24 and SPC25) point in the direction of the microtubule plus-end (**Figure 1**B-D). Thus, the structure explains how HEC1 loop-mediated oligomerization stabilizes kinetochore-microtubule attachment (*5, 17, 28, 29*).

In an unanticipated feature, when neighboring protofilaments are instead considered, the longitudinal stacks of Ndc80Cs alternate with protofilaments occupied by the cryo-EM density corresponding to microtubule-binding domain (MTBD) of SKA1 (SKA1^MTBD^, *orange)*. This occupies the interdimer α/β-tubulin interface, thus leaving the intradimer interface vacant and alternating positions along the continuous Ndc80C oligomer (**Figure 1**B-C).

The SKA1^MTBD^ functionally cooperates with Ndc80C (*7, 30*). It has been previously shown to bind preferentially, but not exclusively, to curved protofilaments at the microtubule end (*7, 22*), but its interaction with microtubules has not yet been structurally elucidated. In their reciprocal arrangement on neighboring protofilaments, SKA1^MTBD^ and the HEC1 CH domain (HEC1^CH^) surround the negatively charged C-terminal tail of α-tubulin (α^E-hook^, *brown)*, sandwiching it through several contacts (**Figure 1**). At the alternating position, left vacant by SKA1, the HEC1^CH^ interacts with the β-tubulin E-hook (β^E-hook^, *red)*.

The interaction of SkaC and Ndc80C is mediated by the C-terminal region of SKA3 *(pink*, SKA3^C-Term^), which binds the HEC1:NUF2 coiled-coil (*4, 5, 16, 17*). Density for SKA3^C-Term^ is clearly visible in proximity of the kink in the HEC1:NUF2 coiled-coil (CC) (**Figure 1**B-C). SkaC forms a dimer of trimers through N-terminal helical segment of each subunit (*31*), but these multimerization modules were invisible in our density, likely because they are connected with SKA1^MTBD^ and SKA3^C-Term^ by flexible linkers and don’t dock on microtubules or Ndc80C (*31*). As a consequence, although SKA1^MTBD^ occupies both protofilaments siding the Ndc80C longitudinal oligomer, it is unclear how any two copies of SKA1^MTBD^ are linked in the SkaC dimer. Collectively, however, the structure provides a comprehensive view of interactions stabilizing the Ndc80C:SkaC assembly on microtubules.

### Stabilization of the longitudinal stacks of Ndc80C

The visible segments of Ndc80C correspond to regions previously implicated in microtubule binding and to the fragments that connect these regions. These include the N-terminal tail of HEC1, a phosphorylation target of Aurora family kinases previously implicated both in microtubule binding and in the interaction of neighboring Ndc80Cs on microtubules (*13-15, 19, 27, 32*); the CH domains of HEC1 and NUF2, also previously implicated in microtubule binding (*15, 33*); and the loop region, previously suggested to mediate lateral interactions of Ndc80 complexes (*29, 34*) (**Figure 2A-E**). All these elements contribute to the stabilization of the Ndc80C stacks.

**Fig. 2.**
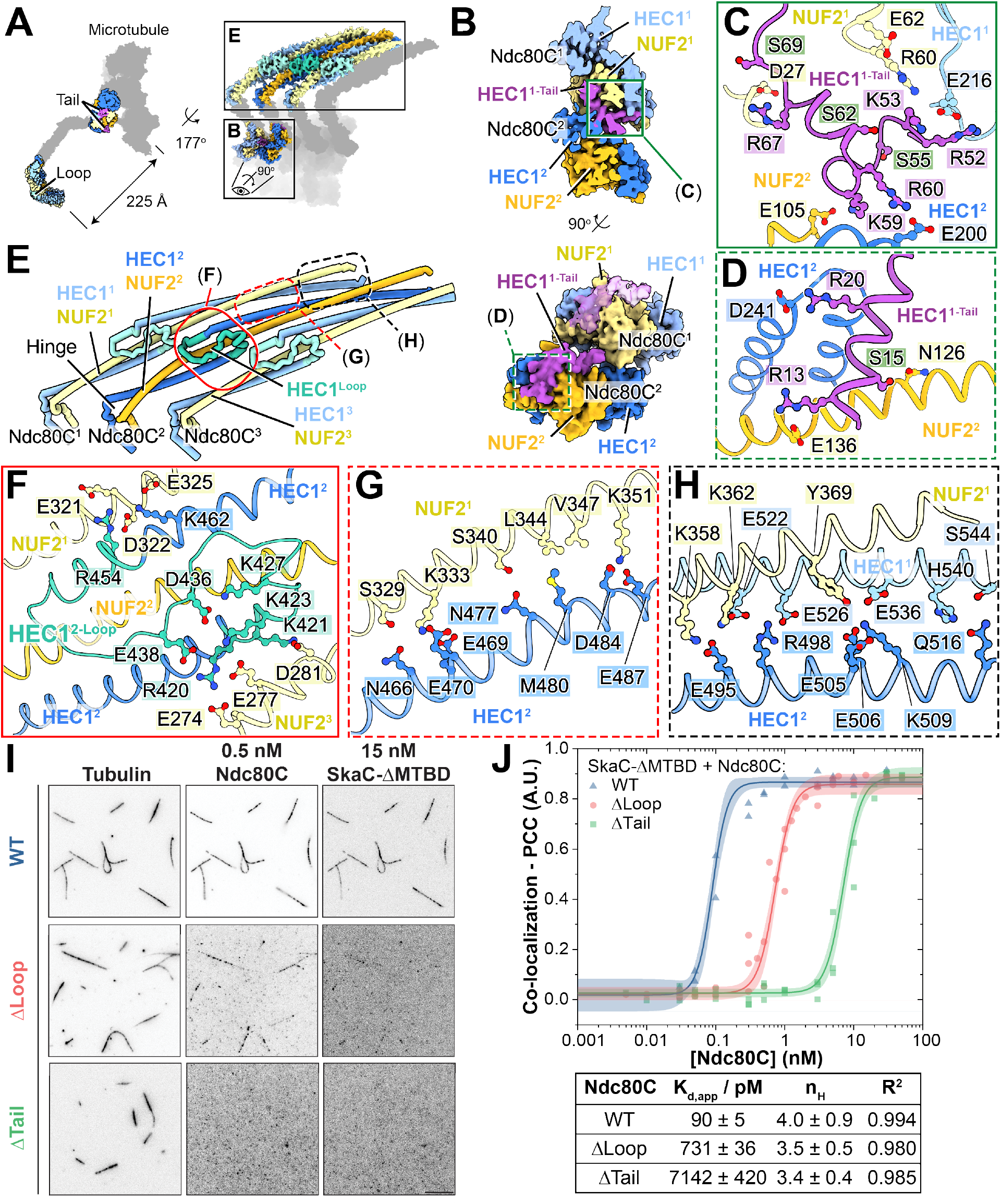
Longitudinal Ndc80C stacks enable high-affinity SkaC binding. **(A)** Cryo-EM density maps of Ndc80C:SkaC bound to microtubules, illustrating the spatial organization of HEC1 N-terminal tail and loop interfaces that mediate Ndc80C oligomerization. **(B)** Cryo-EM density maps highlighting how the HEC1 N-terminal tail of Ndc80C protomer 1 (HEC1^1-tail^, purple) bridges CH domains of two adjacent Ndc80C protomers (Ndc80C^1^ and Ndc80C^2^). (**C and D**) HEC1^1-tail^ bridging Ndc80C^1^ and Ndc80C^2^. Aurora phosphorylation sites in cyan. (**E**) HEC1^loop^-mediated Ndc80C stacks. The HEC1 loop (cyan) bridges neighboring NUF2 coiled-coils (yellow). (**F**) HEC1^loop^ is a switchback, bridging two adjacent NUF2 coiled-coils. (**G** and **H**) Loop-mediated longitudinal contacts at the CCs. (**I**) TIRF microscopy images of Taxol-stabilized microtubules incubated with 15 nM SkaC-ΔMTBD and 0.5 nM of WT, ΔLoop, or ΔTail Ndc80C. Inverted grayscale images were contrast-adjusted individually. Bar: 10 µm. **(J)** Quantification of Ndc80C and SkaC-ΔMTBD colocalization. Pearson correlation coefficients (PCC) at varying concentrations of WT (triangles, n = 2), ΔLoop (circles, n = 3) or ΔTail (squares, n = 3) Ndc80C. Fits to the Hill equation (solid lines) and 95% confidence intervals (shaded) are shown. The apparent dissociation constants *(K*_*d,app*_), Hill coefficients *(n*_*H*_) and *R*^*2*^ derived from the fits are indicated.

The N-terminal tail of HEC1 bridges the CH domains of HEC1 and NUF2 between adjacent Ndc80Cs (indicated as HEC1^1^, HEC1^2^, NUF2^1^, and NUF2^2^ in **Figure 2**B) with hydrophobic and charged-based interactions, as also recently described (*27*). Previously characterized HEC1 tail phosphorylation at S15, S55, S62 and S69 is predicted to interfere with the stabilization of Ndc80C stacks by the HEC1 N-terminal tail (**Figure 2**A-D and **Figure S3**A-B) (*15, 32, 35, 36*). The HEC1 loop (**Figure 2**A,E and **Figure S3**C-E) consists of a structural switchback of the HEC1 subunit, whose chain reverses direction by 180º around residue 426, followed by another inversion ∼20 residues downstream. This double-turn allows the HEC1 chain to resume its original trajectory and reestablish the coiled-coil interaction with NUF2 (**Figure 2**E).

The HEC1 loops contribute a second crucial longitudinal contact between Ndc80Cs, ∼22 nm away from the microtubule lattice where the HEC1 N-terminal tail is located (**Figure 2**A). They line up to form a continuous array along the longitudinal stacks, connecting the distal end of a preceding switchback with the proximal base of the following HEC1 (**Figure 2**E). The stacked arrangement of loops imposes a defined lateral register on the HEC1:NUF2 coiled-coils of subsequent Ndc80Cs, creating a dense bundle (**Figure 2**E). This arrangement is stabilized by several distinct contacts between subsequent Ndc80Cs (closeups in **Figure 2**F-H and **Figure S3**C-E). Those created by the loop itself provide an excellent account of a previous mutational analysis, where alanine mutations in distinct short segments, also comprising R420, K421, K423, K427, D436, E438, and R454, prevented chromosome alignment and activated the spindle assembly checkpoint (*29*). These residues are now shown to mediate or stabilize ionic interactions of the HEC1 loop with NUF2 chains of neighboring Ndc80Cs (**Figure 2**F and **Figure S3**F-H). Downstream of the loop, towards the HEC1 and NUF2 C-termini, several additional ionic contacts between neighboring coiled-coils are observed (**Figure 2**G-H and **Figure S3**I-J). Thus, the HEC1 loop stabilizes the stacking of neighboring Ndc80Cs complexes with a defined long-range register, creating a continuous interface that promotes multiple new lateral ionic interactions between HEC1 and NUF2 in distinct complexes.

In vitro, SkaC binds Ndc80C monomer or its loopless version (Ndc80-ΔLoop) indistinguishably, when Ndc80C does not further oligomerize in solution ((*5*); see also **Figure S4**F). In vivo, however, kinetochore localization of SkaC requires the loop and positively charged residues of the HEC1 N-terminal tail (*17, 18, 28*). This suggests that microtubule-guided Ndc80C oligomerization further promotes SKA3 recruitment to the attached kinetochores. To directly test this, we used total internal reflection fluorescence (TIRF) microscopy to visualize the binding of Ndc80C, Ndc80C-ΔTail, or Ndc80C-ΔLoop to microtubules, and monitored the concomitant binding of a SkaC construct lacking the MTBD (SkaC-ΔMTBD), whose interaction with microtubules is therefore entirely dependent on an interaction with the Ndc80C (**Figure 2**I). By titrating Ndc80C, Ndc80C-ΔTail or Ndc80C-ΔLoop in this experiment, we found that SkaC-ΔMTBD colocalized considerably less efficiently with Ndc80C-ΔTail and Ndc80C-ΔLoop (**Figure 2**J). Cryo-EM visualization confirmed that wildtype Ndc80C forms highly organized arrays on microtubules, whereas decoration with Ndc80C-ΔLoop resulted in an entirely disorganized appearance (**Figure S3**K). These observations support the idea that SkaC binds preferentially to an ordered oligomeric form of Ndc80C on microtubules.

### The SKA3 binding site on Ndc80C

Binding of SkaC to Ndc80C requires phosphorylation of the SKA3 C-terminus (on T358 and T360) by CDK1 (*5, 16, 17*). The molecular basis of this interaction has remained unknown, but our structure identifies the SKA3 fragment (residues 356 and 391) comprising phosphorylated pT358 and pT360 bound to Ndc80C. The SKA3 fragment adopts a hairpin, whose tip (residues 367-370) binds against the positively charged Ndc80C kink centered on NUF2-P173 and HEC1-S295 (**Figure 3**A-C and **Figure S4**A). The phosphates on pT358 and pT360 appear to stabilize the SKA3 chain while also promoting direct interactions with NUF2 and HEC1 (**Figure 3**C). AlphaFold2 (AF2) structural modelling (*37, 38*) also predicted a similar binding configuration (**Figure S4**B), but differing from our cryo-EM model on how the Ndc80C kink interacts with SKA3 (**Figure S4**C-D).

**Fig. 3.**
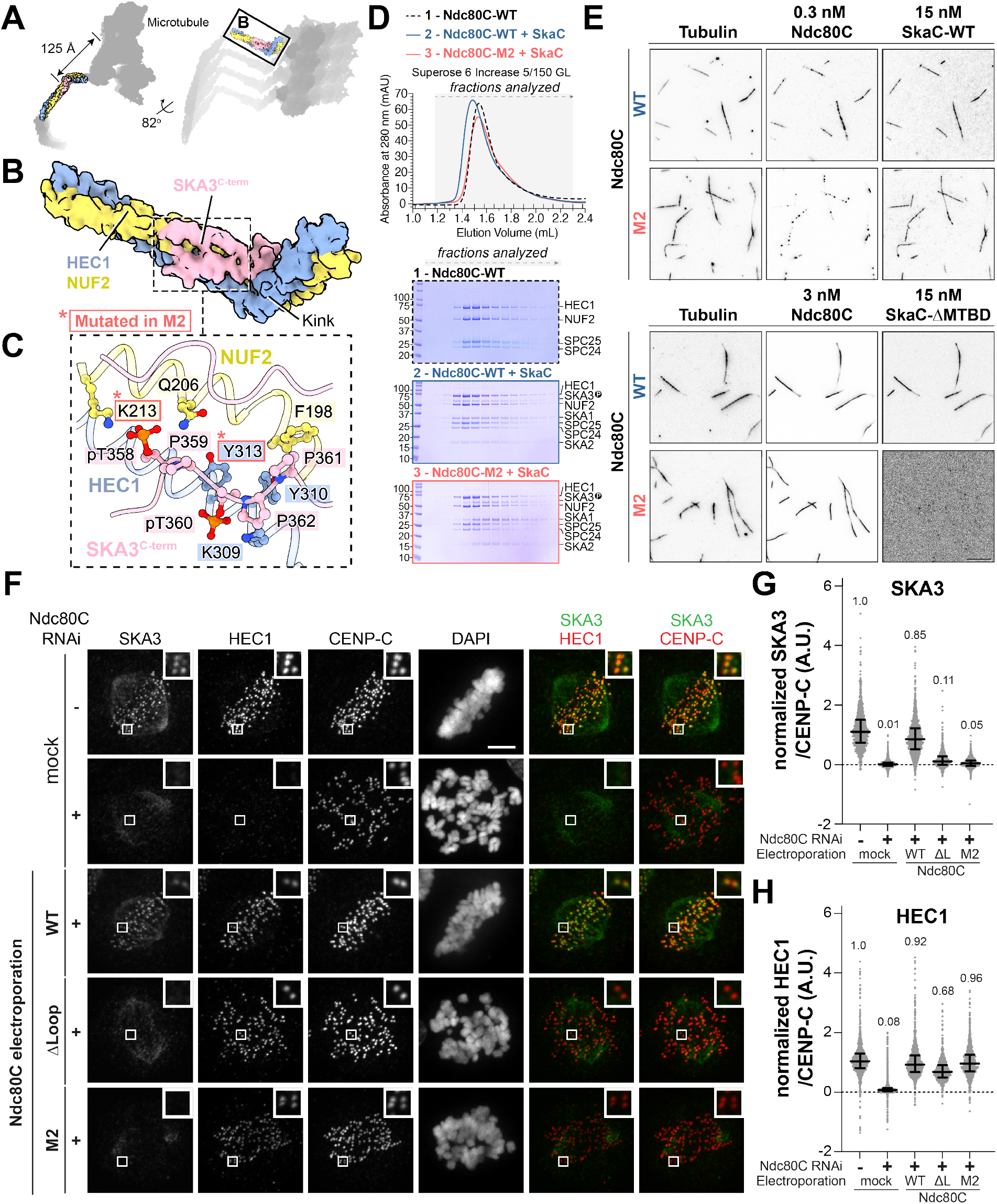
Molecular basis of the CDK1-mediated SKA3 recruitment to the Ndc80C kink. (**A** and **B**) Cryo-EM density maps highlighting binding of SKA3^C-term^ to Ndc80C kink. **(C)** Binding interface of Ndc80C:SKA3 covering CDK1:Cyclin-B phosphorylation sites (pT358/pT360). Residues mutated in Ndc80C-M2 (HEC1 Y313A, NUF2 K213Q) are framed in red. (**D**) Size-exclusion chromatography analysis of phosphorylated SkaC interaction with Ndc80C-WT or -M2. Stable complex is formed with Ndc80C-WT, but not with -M2. (**E**) TIRF microscopy of Taxol-stabilized microtubules decorated with 0.3 nM (top) or 3 nM (bottom) Ndc80C-WT or Ndc80C-M2 with 15 nM SkaC-WT or -ΔMTBD, respectively. SkaC-ΔMTBD colocalizes with Ndc80C-WT but not with Ndc80C-M2. Inverted grayscale images were contrast-adjusted individually to reveal weak binding. Bar: 10 µm. (**F**) Immunofluorescence of HeLa cells untreated or treated with Ndc80C RNAi and electroporated with mock, Ndc80C-WT, -ΔLoop (-ΔL), or -M2. CENP-C marks kinetochores. Bar: 5 µm. (**G** and **H**) Quantification of kinetochore levels of endogenous SKA3 (**G**) and endogenous or electroporated HEC1 (**H**) normalized to endogenous levels (no RNAi). Number of kinetochores analyzed from three independent experiments: n = 2669 (control), 2773 (RNAi), 2508 (WT), 1563 (ΔL), 2506 (M2). Median, and interquartile range of normalized single kinetochore intensities are also shown.

To functionally validate the SkaC:Ndc80C interaction interface, we introduced point mutations in highly conserved HEC1 and NUF2 residues (**Figure S4**E) to test their significance in stabilizing the observed conformation of the SKA3 hairpin. We generated an Ndc80C-M1 mutant (HEC1^L299A^-NUF2^F180A^) guided by the AF2 model; and Ndc80C-M2 (NDC80^Y313A^-NUF2^K213Q^) predicted on the basis of the cryo-EM structure (**Figure S4**C-D). We also combined these mutations in Ndc80C-M3. Size-exclusion chromatography (SEC) analysis showed that wild-type and M1, but not M2 and M3, interacted with SkaC (**Figure 3**D and Figure **S4**F), validating our cryo-EM structure model of the phospho-dependent Ska3C:Ndc80C interface.

Disruption of the SkaC-binding interface in Ndc80C-M2 was further demonstrated in TIRF experiments with microtubules (**Figure 3**E). We have previously shown that the isolated, monomeric Ndc80C exhibits unstable binding to microtubules at ≤ 0.6 nM but it can cluster and stably bind to microtubules at higher concentrations (*27, 29*). In the presence of 15 nM SkaC, Ndc80C exhibits stable clustered binding to microtubules at 0.3 nM *(upper panel)*. Under the same condition, Ndc80C-M2 bound microtubules less effectively, suggesting that SkaC facilitates Ndc80C oligomerization on microtubules. Microtubule decoration by Ndc80C-WT or Ndc80C-M2 was comparable at 3 nM in the presence of SkaC-ΔMTBD, whose colocalization with microtubules is expected to require the interaction of SKA3 with Ndc80C *(lower panel)*. However, SkaC-ΔMTBD bound Ndc80C-WT normally, but was completely unable to bind Ndc80C-M2 (**Figure 3**E). This result further validates in vitro the Ndc80C binding surface involved in SKA3 binding.

To investigate whether mutations also prevent kinetochore recruitment in vivo, we depleted Ndc80C by RNAi in HeLa cells. We then replaced it with wild-type or recombinant Ndc80 complexes by protein electroporation (*39*). Depletion of Ndc80C prevented SkaC recruitment to kinetochores, but re-introduction of Ndc80C restored SkaC levels (**Figure 3**F-H). As previously shown (*28, 29*), Ndc80C-ΔLoop was unable to recruit SkaC to kinetochores. In line with our in vitro interaction results, Ndc80C-M2 and Ndc80C-M3, but not Ndc80C-M1, also failed to recruit SkaC, to establish cold-resistant kinetochore-microtubule attachments, and to achieve metaphase chromosome alignment (**Figure 3**F-H and **Figure S5**A-F). Collectively, these experiments validate the interaction mechanism of SKA3 with Ndc80C and show that it is essential for stable bipolar kinetochore-microtubule attachment.

### SkaC bridges tubulin dimers and surrounds α-tubulin E-hook with Ndc80C

SKA1^MTBD^ consists of a modified winged-helix fold in which a central α3-helix connects two small helical bundles. This arrangement gives the domain an elongated, parallelepiped-like shape (*7, 30*) (**Figure 1**D). Being positioned at the interdimer interface of the protofilament to which it binds predominantly (PF1; **Figure 4**A-B) and making contacts with the neighboring protofilament largely occupied by Ndc80C (PF2), SKA1^MTBD^ effectively sits at the intersection of four α/β-tubulin dimers. We speculate that binding at the interdimer interface may stabilize terminal tubulin dimers at the microtubule plus-end, which presents free β-tubulin, where attachment and dissociation of α-tubulin of the α/β-dimer respectively leads to depolymerization and polymerization of microtubules. This might explain why presence of SkaC decreases the rate of microtubule depolymerization from the plus-end (*5*).

**Fig. 4.**
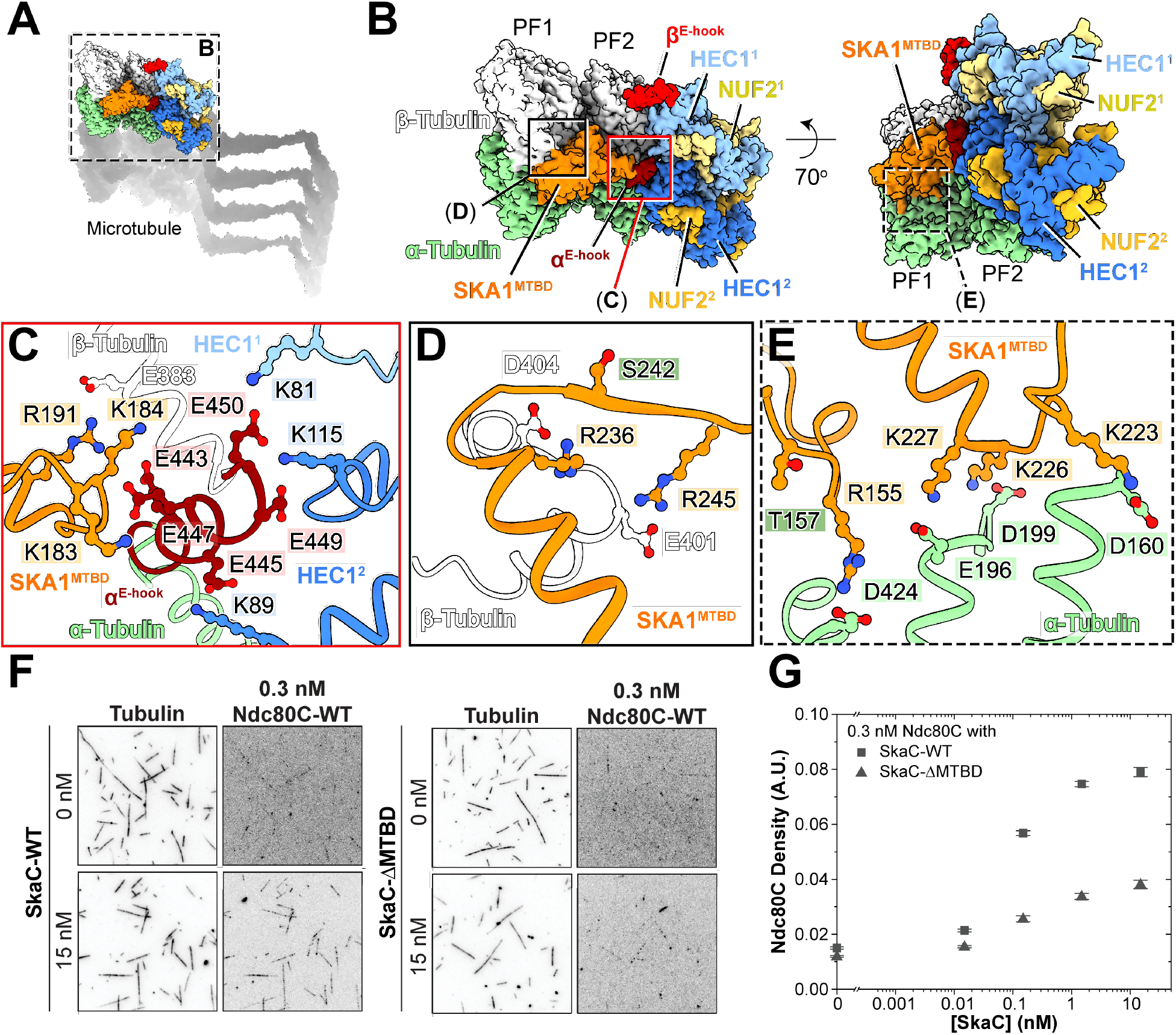
SKA1 promotes cooperative microtubule-binding with Ndc80C. (**A**) SKA1^MTBD^ (orange) location on Ndc80C:SkaC assembly on the microtubule. (**B**) Cryo-EM density maps highlighting SKA1^MTBD^ located at the longitudinal inter-tubulin dimer interfaces of PF1 and PF2. SKA1^MTBD^ also laterally contacts with α-tubulin E-hook (α^E-hook^) of PF2, which is simultaneously bound by two CH domains of Ndc80C (HEC1^1^ and HEC1^2^) on PF2. **(C)** Indirect tethering of SKA1^MTBD^ to Ndc80C stacks via the α^E-hook^ of PF2. (**D**) Interface between SKA1^MTBD^ and β-tubulin of PF1. (**E**) Interface between SKA1^MTBD^ and α-tubulin of PF1. (**F**) TIRF microscopy of Taxol-stabilized microtubules decorated with low density (0.3 nM) Ndc80C-WT in the presence or absence of 15 nM non-fluorescent SkaC-WT or SkaC-ΔMTBD. Inverted grayscale images were contrast-adjusted individually to reveal weak binding. Bar: 10 µm. **(G)** Quantification of the Ndc80C fluorescence signal on microtubules at a fixed concentration of 0.3 nM with varying concentrations of SkaC-WT or SkaC-ΔMTBD.

The SKA1^MTBD^ binds the α-tubulin E-hook, which is simultaneously bound also by two HEC1^CH^ domains of neighboring Ndc80C protomers. This stabilizes the α^E-hook^ to the point that its density is clearly visible (**Figure 4**C and **Figure S6**A). The interaction is mediated by positively charged residues of SKA1^MTBD^ (K183, K184, K185, R191; **Figure S6**B) previously implicated in microtubule binding (*7, 30*). On the two HEC1^CH^ involved, the interaction with the E-hook engages K81 and K115 from one protomer, and K89 from another protomer (**Figure 4**C and **Figure S6**B), also previously implicated in microtubule binding (*15*). Thus, there are no direct contacts between SKA1 and Ndc80C, but the encirclement of the α-tubulin E-hook creates an integrated interface that likely contributes to generating a cooperative interaction of SkaC and Ndc80C on microtubules.

On the other adjacent protofilament, SKA1^MTBD^ bridges the inter-tubulin dimer interface by interacting with both α- and β-tubulin. Here, SKA1^MTBD^ wedges at one end between α- and β-tubulin (**Figure 4**D-E), where positively charged residues including R236/R245 and R155/K223/K226/K227 interact with acidic residues in α- and β-tubulin, including E401 and D404 of β-tubulin, and D160, E196, D199, and D424 of α-tubulin. These residues have been previously identified for a function in microtubule binding (*7, 30*) (**Figure 4**D-E and **Figure S6**C-D).

We used TIRF microscopy to assess whether SKA1^MTBD^ stabilizes the interaction of Ndc80C with microtubules. At subnanomolar concentrations of Ndc80C, addition of SkaC (at 15 nM) strongly increased the decoration of microtubules by Ndc80C (**Figure 4**F-G). Conversely, there was only weak enhancement of Ndc80C binding when SkaC-ΔMTBD was used in place of SkaC wild-type. These results are consistent with the cooperative microtubule-binding of Ndc80C and SkaC, where microtubule-guided Ndc80C oligomerization promotes SkaC binding to Ndc80C (**Figure 2**I-J), SkaC enhances the microtubule-binding activity of Ndc80C, in agreement with the structural work.

## Discussion

The structural and functional work presented here reveals how Ndc80C and SkaC bind to each other and create a cooperative and controlled interaction with microtubules. Our work provides a systematic and comprehensive perspective on the role of most previously identified functional moieties of Ndc80C and SkaC.

The kinetochore is built hierarchically on a specialized chromosome structure, the centromere. Linkages from centromere-proximal kinetochore subunits, including CENP-C and CENP-T in humans, assemble the outer kinetochore under defined stoichiometries. Biochemical and imaging analyses indicate that only a limited number of Ndc80Cs and SkaCs populate each kinetochore microtubule-binding site (*40*). As human CENP-C and CENP-T have been shown to recruit one and three Ndc80 complexes, respectively (*1, 41*), we surmise that each of these functional kinetochore unit can recruit four Ndc80Cs and two SkaC dimers. Further dimerization of CCAN around the specialized centromeric nucleosome may double this number.

How this minimal machinery ensures stable, high-fidelity segregation remains a key unresolved question. While future work will likely visualize these interactions in situ at physiological density, current high-resolution structural analyses rely on fully decorated microtubules in vitro. Despite this limitation, our work provides a clear answer to the initial question. Specifically, the observation that Ndc80C occupies alternating protofilaments (i.e. that it does not form lateral interactions with Ndc80Cs on neighboring protofilaments) and the detailed mechanisms of longitudinal stacking we have described, strongly indicate that Ndc80Cs organize themselves in longitudinal arrays at kinetochores.

We reveal how the HEC1 loop bundles the kinetochore-proximal stems of HEC1:NUF2 coiled-coils (segments downstream of the hinge of HEC1 and NUF2 towards the C-termini of these proteins). As the loop lacks sites for post-translational modifications, we propose it may play a constitutive role in bundling the stems of Ndc80Cs that extend from the kinetochore even before biorientation. Conversely, loop-mediated bundling is essential, albeit not sufficient, for a tight interaction of Ndc80C with SkaC. Single Ndc80C-ΔLoop mutant tetramers bind to SkaC in vitro indistinguishably from Ndc80C-WT at micromolar concentrations (*5*) (**Figure S4**F). Conversely, Ndc80C-ΔLoop fails to recruit SkaC in vivo (**Figure 3**F-H **and Figure S5**A), indicating that the affinity observed in vitro with single Ndc80C tetramers in solution is insufficient for robust kinetochore localization. This is confirmed by our TIRF measurements, which reveal a striking difference in the context of the microtubule lattice: while SkaC binds wild-type Ndc80C with high avidity, this affinity drops nearly 8-fold for the loop-deficient mutant. We speculate that Ndc80C stacking through the loop may contribute to basal recruitment of SkaC, and that the associated protein phosphatase 1 (PP1) activity contributes to the initial dephosphorylation of the N-terminal tail (*9, 42*).

Together, Ndc80C bundling, SKA3 binding to Ndc80C, and the sandwiching of the α-tubulin E-Hook create a continuous binding interface across neighboring protofilaments. This binding mode suggests a compelling mechanism for kinetochore-microtubule stabilization: progressive dephosphorylation of the HEC1 N-terminal tail promotes longitudinal stacking of Ndc80C, generating an ideal high-avidity landing array for SkaC and potentially for other proteins, like the Astrin-SKAP complex (*43, 44*), that stabilize the kinetochore-microtubule interface to license chromosome separation and segregation to the daughter cells.

Since its discovery, animal SkaC has been considered a functional ortholog of yeast Dam1 complex (Dam1C) despite the apparent lack of sequence and structural homologies between the two complexes (*11, 30, 31*). Our study illustrates their clear structural differences. The cryo-EM structure of Ndc80C-Dam1C on microtubules showed that 16 protomers of 10 subunit Dam1C form a ring to encircle a microtubule. The Dam1C^ring^ is positioned toward the microtubule plus-end relative to the CH domains of the Ndc80C (*45*). Unlike human Ndc80C, yeast Ndc80C does not longitudinally oligomerize over a microtubule protofilament, but it laterally clusters via Dam1C^ring^ interactions. Whereas Ska1^MTBD^ bridges longitudinal and lateral α/β-tubulin dimer interactions, Dam1C^ring^ does not exhibit a clear direct contact to the microtubule surface. The Dam1C^ring^ may slide near dynamic microtubule ends, but is hooked on the microtubule surface by Ndc80C. Conversely, cooperative multimeric Ndc80C-SkaC assembly generates an oligomeric Ndc80C sleeve that is coupled with Ska1 that binds preferentially curved microtubule protofilaments.

We propose a model for the progressive maturation of kinetochore attachments (**Figure 5**). Initially, Aurora B-mediated phosphorylation of the HEC1 N-terminal tails restricts the kinetochore to weak, transient interactions to microtubules. As HEC1 is progressively dephosphorylated, the formation of Ndc80C stacks generates a high-avidity platform that robustly recruits SkaC. On mature attachment, the cooperative assembly of Ndc80C and SkaC enables microtubule end-tracking and load bearing by interacting with the straight lattice and curved protofilaments at the same time.

**Fig. 5.**
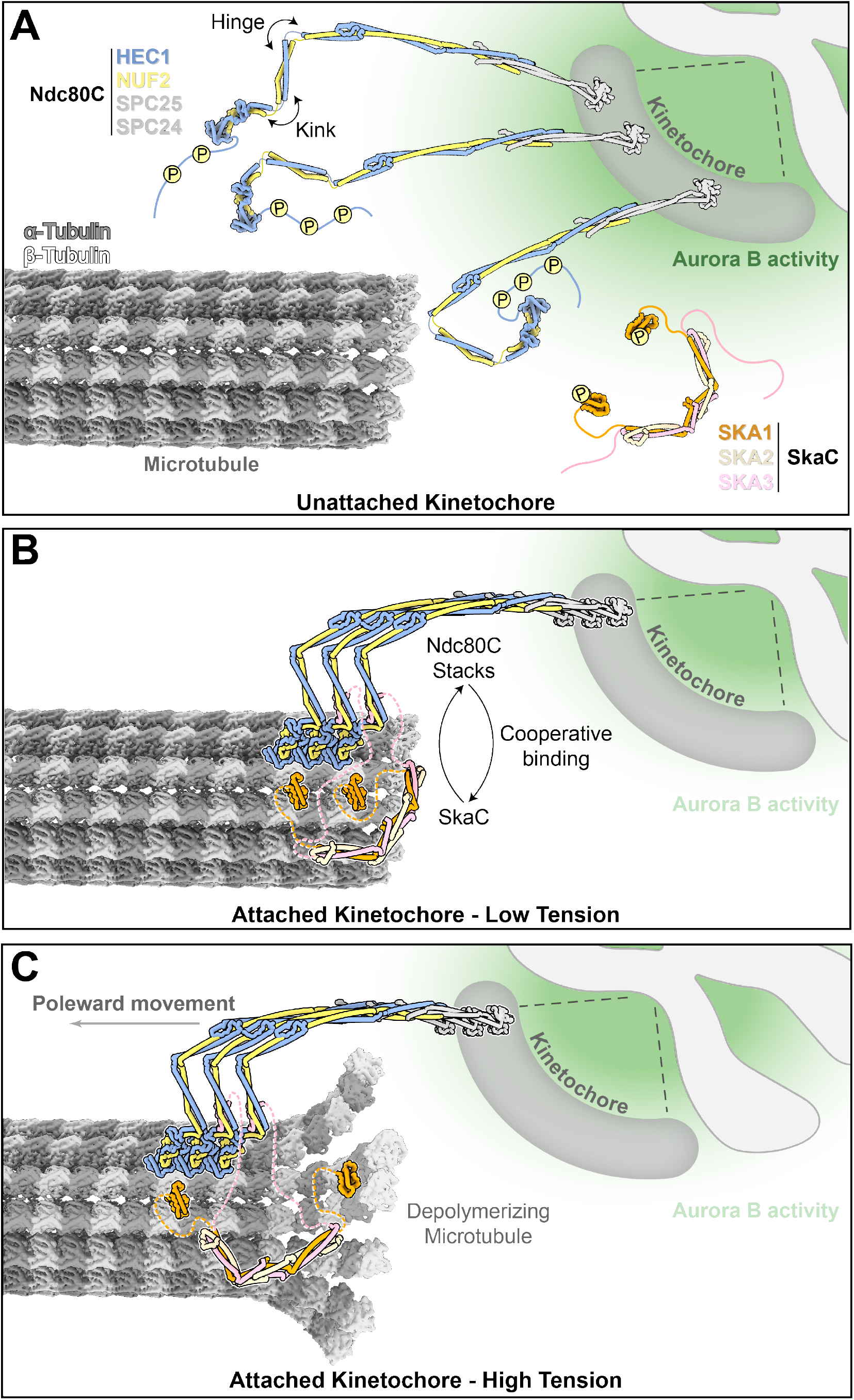
Model for establishing stable end-on kinetochore-microtubule attachments by cooperative assembly of Ndc80C and SkaC. (**A**) Unattached kinetochore, with HEC1 tails and SKA1 phosphorylated by Aurora B (activity indicated by the green gradient). (**B**) Under low-tension conditions Ndc80C can form longitudinal clusters on the microtubule lattice. These clusters allow for cooperative binding of SkaC, which in turn promotes further Ndc80C oligomerization. (**C**) During microtubule depolymerization, stacked Ndc80C assemblies move with the lattice as protofilaments curl, while SkaC preferentially engages curved protofilaments, facilitating kinetochore end tracking. While Ndc80C oligomers offer multiple microtubule-binding sites that enable tracking of dynamic microtubules, SkaC staples both longitudinal and lateral inter-α/β-dimer interfaces to support load-bearing attachment.

## Acknowledgments

We thank P.J. Huis in’t Veld, I. Vetter, and N. Raam Sankar for contributions to the initial phase of this project, M. Terbeck and S. Wohlgemuth for protein productions, R. Gong for advice on cryo-EM analysis, and the help of the High-Performance Computing Resource Center at the Rockefeller University.

## Funding

National Institutes of Health grant R35GM132111 (HF)

European Research Council (ERC) Synergy Grant 951430 (BIOMECANET) (AM)

DGF’s Collaborative Research Centre 1430 “Molecular Mechanisms of Cell State Transitions” (AM).

## Author contributions

Conceptualization: YN, DM, SG, JM, HF, AM

Methodology: YN, DM, SG, JM, DP, OH

Investigation: YN, DM, SG, JM

Visualization: YN, DM, SG, JM

Funding acquisition: HF, AM

Project administration: HF, AM

Supervision: HF, AM

Writing – original draft: YN, DM, SG, JM, HF, AM

Writing – review & editing: YN, DM, SG, JM, HF, AM

## Data and materials availability

Data and material are available from the corresponding authors as applicable. Cryo-EM maps and coordinates have been deposited in the EMDB and wwPDB, respectively, with accession numbers EMD-74572 and PDB 9ZQO; EMD-74573 and PDB 9ZQP (SKA1, HEC1 tail and microtubule interface).

## Supplementary Materials

Materials and Methods

Figs. S1 to S6

Tables S1

References (*47–70*)

