## Supplementary Material for "Molecular basis of cooperative assembly of the Ndc80-Ska kinetochore complex on microtubules"

### Molecular basis of cooperative Ndc80-Ska complex assembly on microtubules for kinetochore-microtubule attachment

**The PDF file includes:**

Materials and Methods
Figs. S1 to S6
Table S1
References 47-69

**Materials and Methods**

Cloning and mutagenesis

Ndc80C-M1, -M2 and -M3 mutants in this study were generated on pLIB vectors containing either NUF2-WT or HEC1-WT by PCR-based site-directed mutagenesis with the following primers: NUF2 F180A 5'-GAAGAGCAAGAAGAGGCAAAGCAGCTTTCAG-3', NUF2 K213Q 5'-GGGAAATTCCCAAAAGCAATCAAATATTTCAGAG-3', HEC1 L299A 5'-GAGTCGTTGAGAAAAGCAAAGGCTTCCTTAC-3', HEC1 Y313A 5'- CAAAAGTATCAG GCAGCTATGAGCAATTTGGAGTC-3'). Mutagenesis was confirmed by Sanger sequencing.

Expression and purification of Ndc80C

Expression cassettes from pLIB vectors containing HEC1, NUF2, SPC25$^{SORT-6xHIS}$, and SPC24 were combined on a pBIG1 vector using Gibson assembly as described (*46*). Baculoviruses were generated in *Sf9* insect cells and used for protein expression in *Tnao38* insect cells. Between 60 and 72 hr post-infection, cells were washed once in PBS and stored at -80 °C. All subsequent steps were performed on ice or at 4 °C.

Cells were thawed and resuspended in lysis buffer (20 mM HEPES pH 8.0, 150 mM NaCl, 10 % v/v glycerol, 1 mM TCEP, 20 mM imidazole) supplemented with 1 mM PMSF, and protease-inhibitor mix HP Plus (Serva), lysed by sonication, and cleared by centrifugation at 98,000 xg for 45 min. The cleared lysate was filtered (0.8 µm) and applied to a 10 or 20 mL HisTrap FF (GE Healthcare) equilibrated in washing buffer (lysis buffer without protease inhibitors). The column was washed with approximately 20 column volumes of washing buffer and bound proteins were eluted with elution buffer (washing buffer containing 300 mM imidazole). Relevant fractions were pooled, diluted 5-fold with buffer A (50 mM HEPES, pH 8.0, 25 mM NaCl, 5 % v/v glycerol, 1 mM EDTA, 2 mM TCEP) and applied to a 25 mL Source 15Q (GE Healthcare) strong anion exchange column equilibrated in buffer A. The column was washed with 3-5 column volumes of buffer A and proteins were eluted with a linear gradient from 25 mM to 300 mM NaCl in 180 mL. Relevant fractions were concentrated in 30 kDa molecular mass cut-off Amicon concentrators (Millipore) and applied to a Superdex 200 16/60 column (GE Healthcare) equilibrated in 20 mM HEPES pH 8.0, 250 mM NaCl, 5 % v/v glycerol, 1 mM TCEP. Relevant fractions were pooled, concentrated, flash-frozen in liquid nitrogen, and stored at -80 °C. The same protocol was used to express and purify the Ndc80C-ΔL mutant (HEC1-Δ430-462) and the following point mutants: M1 (HEC1-L299A, NUF2-F180A), M2 (HEC1-Y313A, NUF2-K213Q), and M3 (combined M1 and M2 mutations).

Expression and purification of phorphorylated SkaC-WT and -ΔMTBD

Expression cassettes from pLIB vectors containing SKA1$^{SORT-6xHIS}$, SKA2, and SKA3 were combined on a pBIG1 vector using Gibson assembly as described (*46*). Baculoviruses were generated in *Sf9* insect cells and used for protein expression in *Tnao38* insect cells. Between 60 and 72 h post-infection, cells were washed once in PBS and stored at -80 °C. All subsequent steps were performed on ice or at 4 °C. Cells were thawed and resuspended in lysis buffer (20 mM HEPES, pH 8.0, 150 mM NaCl, 10 % v/v glycerol, 1 mM TCEP, 20 mM imidazole) supplemented with 1 mM PMSF, and protease-inhibitor mix HP Plus (Serva), lysed by sonication, and cleared by centrifugation at 98,000 x *g* for 45 min. The cleared lysate was applied to 5 mL of cOmplete™ beads (Roche) equilibrated in lysis buffer and incubated at 4 °C overnight for binding. The beads were washed twice with ~30 mL of lysis buffer, and eluted for 1 h at 4 °C in elution buffer (lysis buffer with 300 mM imidazole). The elution fraction was diluted 10-fold with buffer A (20 mM HEPES, pH 8.0, 25 mM NaCl, 5 % v/v glycerol, 1 mM TCEP), filtered (0.8 μm) and applied to a 25 mL Source15Q (GE Healthcare) strong anion exchange column equilibrated in buffer A. The column was washed with 3-5 column volumes of buffer A and proteins were eluted with a linear gradient from 25 mM to 300 mM NaCl in 180 mL. Relevant fractions were concentrated in 30 kDa molecular mass cut-off Amicon concentrators (Millipore) and treated with 75 nM CDK1:CyclinB (with 2 mM ATP and 10 mM $MgCl_2$) overnight at 5 °C or for 4 h at 25 °C. The reaction mix was then applied to a Superdex 200 16/60 column (GE Healthcare) equilibrated in 20 mM HEPES, pH 8.0 200 mM NaCl, 5 % v/v glycerol, 1 mM TCEP. Relevant fractions were pooled, concentrated, flash-frozen in liquid nitrogen, and stored at -80 °C. The SkaC-ΔMTBD variant (SKA1-Δ109-255) was expressed and purified using the identical protocol.

Fluorescent labeling of Ndc80C and SkaC

The calcium-independent Sortase 7M (*47*) was used for the C-terminal conjugation of a synthetic peptide to LPETGG-tagged subunits. Various synthetic peptides (Genscript) were used for this purpose: GGGGC-Alexa Fluor 488 (used for Ndc80C) and GGGGK-TAMRA (used for SkaC). Reactions were performed at 4 °C for 16 h or at 25 °C for 4 h in the respective size-exclusion buffer with a Sortase:target:peptide ratio of approximately 1:10:50 with the target protein in the 10–20 μM range. Excess peptide and Sortase were removed from the fluorescently labeled complexes using size-exclusion chromatography, as described above.

Tubulin and microtubule polymerization

Tubulin was purified from porcine brain extract using the high-molarity PIPES cycling method (*48*). Fluorescent and biotinylated tubulin were prepared using NHS-ester derivatives according to published protocols (*49*). Depolymerized (labeled and unlabeled) tubulin was flash-frozen in liquid nitrogen as single-use aliquots (10 or 20 μL) and stored at -80 °C.

For cryo-EM, microtubules were polymerized by diluting unlabeled tubulin (125 μM) to 50 μM in polymerization buffer (25 mM MES pH 6.8, 70 mM NaCl, 1 mM $MgCl_2$, 1 mM EGTA) supplemented with 3 mM GTP. The reaction was incubated at 37 °C for 90 min and stabilized by addition of Taxol to a final concentration of 200 μM. Microtubules were pelleted by centrifugation at 17,000 x *g* for 8 min at 22 °C, resuspended in polymerization buffer containing 3 mM GTP and 20 μM Taxol, and pelleted again under the same conditions. After discarding the supernatant, microtubules were resuspended in binding buffer (30 mM HEPES KOH pH 7.2, 5 mM $MgSO_4$, 1 mM EGTA, 1 mM DTT, 5 μM Taxol). Digitonin was added to a final concentration of 0.05 % immediately prior to grid preparation.

Microtubules for TIRF experiments were polymerized by mixing unlabeled tubulin, Atto643-labeled and biotinylated tubulin (8:1:1 molar ratio) in BRB80 (80 mM K-PIPES pH 6.8, 1 mM $MgCl_2$, 1 mM EGTA) supplemented with 1 mM GTP, to a final tubulin concentration of 50 μM. The mixture was incubated at 37 °C overnight and stabilized with 50 μM Taxol. Microtubules were centrifuged at 17,000 x *g* for 8 min at 22 °C, resuspended in double the initial volume of fresh BRB80 containing 40 μM Taxol (BRB80T40), and centrifuged again. The final pellet was resuspended in BRB80T40 the same way as before and stored at room temperature for up to three days.

Cleaning and passivation of coverslips

Glass coverslips were cleaned using a protocol adapted from (*50*). Briefly, coverslips were sequentially washed with water, 5 % Hellmanex III, and ethanol, followed by 10 min of argon/oxygen plasma cleaning. Surfaces were then silanized by incubating for 1 h in 0.05 % dichlorodimethylsilane in heptane (adapted from (*51*)). Finally, coverslips were rinsed with heptane and dried in a 40 °C chamber.

Preparation of flow chambers and TIRF microscopy

Flow chambers were assembled using 6-well bottomless sticky-Slides (ibidi) and silanized coverslips. Neutravidin was adsorbed directly onto the silanized glass, followed by surface passivation with 1 % Pluronic F-127 and 1 mg/mL Blocker-Casein (Thermo Scientific) in BRB80. Taxol-stabilized biotinylated microtubules were diluted in BRB80 supplemented with 10 µM Taxol (BRB80T10) and introduced into the chamber. After a 5 min incubation, unbound microtubules were removed by washing with 3 x100 µL of BRB80T10. Subsequently, proteins of interest were added into the chamber in imaging buffer (BRB80, 1 mg/mL Blocker-Casein, 4 mM DTT, 0.2 mg/mL Catalase, 0.4 mg/mL Glucose Oxidase, 20 mM Glucose). The imaging buffer was pre-cleared by ultracentrifugation (Beckman TLA-120.1) for 20 min at 120,000 rpm prior to the addition of proteins.

TIRF microscopy

Experiments were performed using a Nikon Ti-E inverted microscope with a CFI Apo 100x/1.49 NA oil-immersion objective (Nikon) and an iLas2 ring TIRF module (Visitron Systems). Excitation was achieved using 488 nm, 561 nm and 640 nm lasers. Emission light was directed to either of two Prime 95B sCMOS cameras (Teledyne Photometrics) via a TwinCam beam splitter (Cairn). The splitter contained a T560lpxr (Chroma) dichroic mirror to separate channels, followed by ET525/50m, or ET605/52m and ET700/75m emission filters (Chroma). Images were acquired sequentially (640 nm, 561 nm, then 488 nm) with 100 ms exposure time per channel at room temperature using VisiView software (Visitron Systems). The files were stored as 16bit OME-TIFF with an effective pixel size of 110 nm.

Image analysis and quantifications

Image analysis was performed using custom-written Python 3 software. Backgrounds were corrected by subtracting the median intensity of manual ROIs. Microtubules were segmented from the 640 nm channel using Kuwahara filtering (radius = 3 pixel) and a white top-hat transform (size = 5 pixel), followed by thresholding at the 95$^{th}$ percentile of non-zero pixels. Skeletons were fragmented at junction points (>3 neighbors) to separate crossing filaments. Tracks were manually verified and intensities for each channel were independently extracted by averaging over a 5-pixel width along the filament coordinates.

Co-localization was quantified by calculating the Pearson correlation coefficient (PCC) between the fluorescence intensity profiles of individual microtubules using the SciPy library. To determine the average correlation per experimental condition, PCCs were Fisher $z$-transformed. The mean $z$-score was calculated for each field of view (replicate), and these values were back-transformed to report the mean correlation. Variability is reported as the standard error of the mean (s.e.m.) of the field-of-view averages.

Protein binding was quantified as the fluorescence intensity ratio (Protein/Tubulin) for each pixel along the microtubule track. The median of these pixel-wise ratios was determined for each individual microtubule, pooled for one condition and reported as the mean binding density with the standard error of mean. Data were plotted and fitted using OriginPro 2019b (OriginLab Corporation, Northampton, MA, USA).

Grid preparation and cryo-EM data acquisition

Grids were prepared using a Vitrobot Mark IV (Thermo Fisher Scientific) at 22 °C and 100 % humidity. Taxol-stabilized microtubules (2 μL at 2 μM) were applied to glow-discharged UltrAuFoil R1.2/1.3 holey gold 300-mesh grids (Quantifoil). After a 1 min incubation, 2 μL of Ndc80C (2 μM) was added, followed by another 1 min incubation and the addition of 2 μL of SkaC (2 μM). Following a final 3 min incubation, excess liquid was removed by blotting (3.5 s at blot force −3) and grids were vitrified in liquid ethane. All proteins were diluted in binding buffer (30 mM HEPES pH 7.2, 5 mM $MgSO_4$, 1 mM EGTA, 1 mM DTT, and 5 μM Taxol) supplemented with 0.05 % digitonin.

Cryo-EM data were recorded using the EPU software on a Cs-corrected Titan Krios G2 transmission electron microscope operated at 300 kV and equipped with a field emission gun, at a nominal magnification of 81,000× corresponding to a physical pixel size of 0.88 Å per pixel. Images were collected using a zero-loss slit width of 15 eV on the Bioquantum energy filter (Gatan) and a defocus range from -1.5 to -2.3 μm with a K3 direct electron detector (Gatan) in super-resolution counting mode. A total dose of ~58 electrons per $Å^2$ was distributed over 60 frames. Details on data acquisition and processing are summarized in **Table S1**.

Cryo-EM data processing

Data were processed in CryoSPARC v4.6-4.7 (Structural Biotechnology Inc.) (*52, 53*) using a standardized preprocessing workflow. Motion correction and contrast transfer function (CTF) estimation were performed with CryoSPARC's patch-based algorithms (*54*) on the total of 20,502 movies and a representative motion-corrected micrograph is shown in **Figure S1**B. Particles were selected using CryoSPARC's filament tracer (*55*) based on templates simulated from 2D averages of the helical-averaged Ndc80C$^{bonsai}$-microtubule structure (EMD-5489) and custom parameters (filament diameter: 164 Å; separation distance: 82 Å; lowpass filter: 50 Å; Gaussian blur standard deviation: 0.05 and hysteresis low/high thresholds: 88/93 %, respectively). Selected particles were extracted into ~830 Å boxes with 4× binning and cleaned through four rounds of 2D classification disabling option of re-centering 2D averages, yielding 3,904,045 well-centered microtubule particles (**Figure S1**C). Cleaned particles were subjected to heterogeneous refinement using simulated 3D references of 11-16 protofilament (PF) microtubules (https://github.com/moores-lab/MiRPv2), producing the following distribution: 11PF (12 %), 12PF (23 %), 13PF (48 %), 14PF (8 %), 15PF (5 %), and 16PF (4 %). A total of 1,892,908 13PF particles were selected and refined using non-uniform regularization (*56*) under helical refinement with a Twist of 0° and Rise of 82 Å. The resulting reconstruction shows complete MT decoration, but with mixed features corresponding to poorly resolved CH domains of Ndc80C and SKA1-MTBD density at lower threshold, indicating highly heterogeneous binding modes of the two complexes across PFs (**Figure S1**D). To address this, we first sought to identify homogeneous decoration modes of each complex on individual PFs using a PF-sorting strategy detailed below.

PF sorting stream

The refined 13PF particles, processed using helical parameters of Twist 0° and Rise 82 Å with C1 symmetry, were first subjected to symmetry expansion using the estimated Twist and Rise values of −27.72° and 9.239 Å at n=13 in CryoSPARC, resulting in 14,267,682 particles. Local refinement across the microtubule lattice produced averaged densities without seam alignment, after which signal subtraction was applied to retain only the density for a double-PF as well as their decoration densities. The subtracted double-PF set was then locally refined and classified into five 3D classes without image alignment using PCA-initialized references generated from 10,000 particles per class (unless mentioned, this setting is default for all 3D classification step thereafter) and a focused mask covering the CH domains of HEC1 and NUF2 across the entire double PF. Four distinct classes were obtained, corresponding to the following decoration patterns on the two PFs: (1) Ndc80C CH domains (N) on the first PF and SKA1-MTBD (S) on the second PF (class1, NS class); (2) S on the first PF and N on the second with swapped order (class2, SN class), (3) N on both PFs (class3, NN class), and (4-5) S on both PFs, shifted by 41.63 Å (**Figure S1**D). The NS class (class 1) was selected, re-extracted, reconstructed, and locally refined using a mask covering densities of N and N-bound PF2, as well as its adjacent PF3 and decoration densities that were previously not refined nor classified (**Figure S1**D). The refined particles were then signal subtracted to retain only PF2, PF3 and their decorations, locally refined, and subjected to a similar focused 3D classification approach into four classes, yielding two SS classes (class 1 and 2), one NN class (class 3), and one SN class (class 4). The SN class was selected, reconstructed, and manually rotated in UCSF Chimera (*57*) using the *fitmap* command to align with the previously obtained SN class; the resulting rotation matrix was converted to Euler angles and applied to the particle alignment in CryoSPARC via volume alignment job (**Figure S1**D). The rotated SN particles were merged with the earlier SN class, re-extracted, reconstructed, and signal-subtracted to retain only the PF and decoration densities, followed by local refinement of 3,532,678 particles. The resulting map displayed mixed α- and β-tubulin features (**Figure S1**D), prompting further 3D classification of these double-PF SN-decorated particles using a focus mask covering the entire double-PF without image alignment. Two distinct PF classes were obtained, one shifted by 42.89 Å, both exhibiting clear and distinct density in α-tubulin S9-10 and the Taxol-binding pocket of β-tubulin (**Figure S1**D). These two classes were manually aligned and combined, yielding 2,293,731 corrected double-PF SN-decorated particles, which were then re-centered near the CH domains of HEC1 and NUF2, re-extracted, and reconstructed, producing clear features of SKA1-MTBD and CH domains of Ndc80C. The resulting map revealed SKA1-MTBD bound specifically at the junctions of laterally contacting, homotypic tubulin dimers, while the CH domains of HEC1 and NUF2 bind every tubulin monomer longitudinally, positioning the two complexes on neighboring PFs (**Figure S1**D). To determine which complex resides lateral to the longitudinal Ndc80C array, we generated a focus mask covering both S and N densities as well as the decoration densities on the PF adjacent to N and performed another round of focused 3D classification without image alignment. This yielded two poorly aligned classes (class 2 and 3) and two distinct classes: one showing an additional lateral Ndc80C array (SNN, class 1), and another in which the central PF-bound monomeric array of Ndc80C is flanked by SKA1-MTBDs on both neighboring PFs (SNS, class 4). The SNS class was selected, re-extracted, and locally refined, yielding the corrected SNS-PF class with 917,274 particles. To better determine the Ndc80C:SkaC structures on microtubule, these particles were symmetry expanded with Twist and Rise set to 0° and 176.3 Å so that refinement only covers four copies of CH domains of HEC1, NUF2, SKA1-MTBDs and 3 PFs (each containing 2 tubulin hetero-dimers) (**Figure S2**A). These particles were re-extracted with parameter recenter_using_aligned_shifts option on (referred as re-centered). To remove truly duplicates, these re-extracted particles were applied to a publicly available custom script that identifies particles with identical alignment3D/pose and outputs their UIDs (check_pose_duplicates.ipynb, cutoff=1e-5, https://github.com/cryoem-uoft/cryosparcexamples/blob/main). These potential duplicating particles were manually selected via the *cryosparc.dataset.take* method by UID mapping under the python-based cryosparc-tools library (https://tools.cryosparc.com/api/tools.html). Meanwhile, particles separated within 20 Å were selected and the intersected particles through mapping their UIDs, were considered as true duplicates and removed. This procedure refers to the remove-duplicates procedures in later stage of processing, which yielded 1,926,924 particles. These particles were locally refined, and the resulting reconstruction shows weak but clear features of HEC1 tail domain (purple) and E-hook of both α- and β-tubulin (red) at lower contour level (**Figure S2**A).

SKA3, HEC1-kink and HEC1-loop interfaces in the CC domains of Ndc80C

To resolve the CC domains of Ndc80C, we began with the reconstruction of 1,926,924 particles centered on the CH domains of HEC1 and NUF2. To increase box size, this reconstruction and its corresponding particles were re-centered to the end of the Ndc80C CH domains and re-extracted using a 560-pixel (492.8 Å) box. Local refinement was first performed with maximum alignment resolution limited to 8 Å. The resulting reconstruction showed blurred but continuous density extending from the unique kink region of each Ndc80C molecule at low threshold (**Figure S2**B), along with weaker distal densities corresponding to potential HEC1 Loop interfaces, clearly visible in top-projection plots (**Figure S2**B, red circles). To better resolve these regions, we generated a focus mask covering the CC domains across the entire lattice and performed 3D classification to eliminate poorly aligned particles. Focused 3D classification was carried out using PCA initialization with 111,111 particles, class similarity 0.2, O-EM learning rate init 0.1, and lowpass/target resolutions of 3 Å into five classes. Two poorly aligned classes were removed prior to subsequent local refinement (maximum alignment resolution at 8 Å). This iterative process continued until the Ndc80C CC region displayed a clear shape after local refinement (**Figure S2**B), yielding 515,499 particles. Observing substantial improvement of the CC region across the lattice, we conducted a second round of symmetry expansion focusing on the central 83.55 Å unit (**Figure S2**B). After re-centering, re-extraction, and the remove-duplicates procedure (1,182,547 particles), we generated a refined mask encompassing the clearly resolved CC domain of Ndc80C. This refine mask includes four Ndc80Cs, four SKA1-MTBDs, and three PFs each containing two tubulin heterodimers, which was applied to the subsequent local refinement without alignment-resolution limits. We then performed focused 3D classification using a mask covering Ndc80C and SKA1-MTBD, with hard classification, initial/target resolution at 3 Å, reset and per-particle scale optimization, into 16 classes. Two classes (195,603 particles) exhibited substantially improved reconstruction of Ndc80C and SKA1-MTBD following local refinement (**Figure S2**B). These particles were transferred to RELION4 (*58*) using the *cspar2star.py* script from the pyem program (https://github.com/asarnow/pyem.git), and the metadata fields rlnHelicalTubeId and rlnAnglePsiPrior were removed using *relion_star_handler* command, followed by local refinement with a 4 Å lowpass filter, angular sampling/local search starting at 0.9°, and initial offset range/step of 2 and 1 pixels, respectively. Signal subtraction was then performed in RELION, retaining only densities inside the refinement mask, and the resulting particles were imported back into CryoSPARC for another cycle of local refinement. The aligned and subtracted particles were subjected to local CTF refinement to improve reconstruction. These CTF-refined particles were further processed using reference-based motion correction (RBMC, Bayesian polishing) (*59, 60*) in CryoSPARC, using three parallel jobs in extensive-search mode for parameter fitting; all resulting parameter sets were independently applied, and the motion-corrected particles producing the highest-quality reconstruction were retained (192,646 particles).

These particles were then signal subtracted in RELION using the same procedure and imported back into CryoSPARC for additional local refinement and local CTF refinement. Finally, particles were exported to RELION for reconstruction and one additional round of CTF refinement (*61*), yielding a final reconstruction at 3.97 Å (192,646 particles), which was locally sharpened in CryoSPARC to generate the deposited full map. In this map, the Ndc80C loop interface forms a spacing of ~41.77 Å analogous to that of the tubulin monomer (**Figure S2**C). Mask-corrected FSC curves were computed in CryoSPARC and resolutions reported using the 0.143 criterion (**Figure S2**E). Local resolution estimation was also performed in CryoSPARC (**Figure S2**F).

<u>SKA1, HEC1 tail and microtubule interface</u>

Since the initial reconstruction of 1,926,924 particles showed clear feature for the HEC1 tail domain (purple) and the E-hooks of both α- and β-tubulin (red) at low threshold (**Figure S2**A), we began focused 3D classification using this consensus map. A focus mask covering four Ndc80C molecules, SKA1-MTBD, and the tubulin E-hooks (**Figure S2**B) was applied to generate 10 classes. Two classes (375,726 particles) displayed clear features for SKA1-MTBD, the HEC1 tail, and the tubulin E-hooks; these particles were selected for local refinement, which further improved density in these regions. To improve reconstruction, we performed a second symmetry expansion that reduced the 176.3 Å unit to an 83.45 Å (twofold expansion). The expanded particles were re-centered, re-extracted and applied to the remove-duplicates process, resulting in 638,754 unique particles. These particles underwent local refinement followed by a second round of focused 3D classification using the same mask, with hard classification into 8 classes. Two classes (418,785 particles) showed strong features in these regions of interest. This follows the RELION-based particle-subtraction procedure, retaining only the densities inside the refinement mask, and imported the subtracted particles into CryoSPARC for local CTF refinement and RBMC/Bayesian polishing. The cycle of RELION particle subtraction followed by CryoSPARC local refinement and local CTF refinement was repeated, and the final particles were then exported back to RELION for reconstruction with an additional round of CTF refinement, yielding a 3.27 Å map (418,034 particles), which was sharpened in CryoSPARC to produce the deposited full reconstruction. Mask-corrected FSC curves were calculated in CryoSPARC using the 0.143 criterion (**Figure S2**G), and local resolution estimation was also performed in CryoSPARC (**Figure S2**H).

<u>Model building and refinement</u>

For the SKA1, HEC1 tail, and microtubule interface (map2), the starting models are summarized as follows. Tubulin dimers were taken from a previously reported structure (PDB: 7SGS) (*62*). The CH domains of Ndc80C were generated using AlphaFold3 (AF3) (*63*) with HEC1 (O14777, NDC80_HUMAN, residues 1-279) and NUF2 (Q9BZD4, NUF2_HUMAN, residues 1-173). The SKA1-MTBD (AF-Q96BD8-F1, residues 127–255) was obtained from the AlphaFold Protein Structure Database. These were first rigid-body fitted into map2 using ChimeraX. The HEC1 tail regions spanning residues 1-10 and 25-63 were initially removed, as they were largely outside the resolved density. The fitted models were then combined and subjected to refinement in ISOLDE (*64*) within ChimeraX (*65*), using self-referenced position, torsion and secondary-structure restraints, followed by manual inspection and correction in Coot (*66*). Based on the cryo-EM density, the HEC1 tail was manually extended to include residues 10-34 and 51-63. For the tubulin E-hooks, the cryo-EM map quality was sufficient to manually trace the main chain in a helical conformation (**Figure S6**A, bottom panel). Side-chain assignment using α-tubulin (TBA1B_PIG) and β-tubulin (TBB_PIG) was compatible with the experimental density, and the tubulin molecules were accordingly assigned. The resulting models underwent iterative rounds of refinement in ISOLDE (*64*) and Coot (*66*), and finalized by *phenix.real_sapce_refinement* command (*67-69*) to optimize model geometry and optimal fit into the experimental map. All refinement and validation statistics are summarized in **Table S**1. Model-map resolution was determined in Phenix and reported using the 0.5 criterion (**Figure S2**G). Representative densities for all discussed interactions revealed by this map are shown in **Figure S**3A-B and **Figure S6**A, C-D.

For the SKA3, HEC1-kink and HEC1-Loop interfaces in the CC domains of Ndc80C (map1), the refined model mentioned above was manually docked into the experimental map. E-hook of β-tubulins from the lateral PFs to the central Ndc80C molecules were removed and the remaining model was refined in ISOLDE (*64*) applying the self-referenced torsion and distance restrains. An AF3 predicted model with NDC80_HUMAN (O14777, residues 291-480), NUF2_HUMAN (Q9BZD4, residues 173-310 with pS247) and SKA3_HUMAN (Q8IX90, residues 251-412 with pT358 and pT360) was manually docked into the experimental map by *fitmap* command in ChimeraX (*65*) by selecting HEC1 and NUF2 as the starting model, since it adapts a very similar bending angle near the HEC1 hinge region. For the HEC1-Loop region, AF3 predicted model with NDC80_HUMAN (O14777, residues 357-642), NUF2_HUMAN (Q9BZD4, residues 241-466 with pS247), SPC24_HUMAN and SPC25_HUMAN was manually docked into the experimental map by *fitmap* command in ChimeraX by selecting HEC1 resides 395-475 and NUF2 residues 275-314. These docked models were refined by ISOLDE (*64*) applying the self-referenced torsion, distance and secondary structure constrains, which were manually inspected and connected in Coot (*66*) based on the experimental map. NUF2 pS247 was changed back to serine, while SKA3 pT358 and pT360 were kept, and residues in the Ndc80C CC domains outside the densities were removed. The resulting models underwent iterative rounds of refinement in ISOLDE (*64*) and Coot (*66*), and finalized with *phenix.real_sapce_refinement* command (*67-69*) to optimize model geometry and optimal fit into the experimental map. All refinement and validation statistics are summarized in **Table S1**. Model-map resolution was determined in Phenix and reported using the 0.5 criterion (**Figure S2**E). Representative densities for all discussed interactions revealed in this map are shown in **Figure S3**C-E and **Figure S4**A.

<u>Analytical size-exclusion chromatography</u>

8 μM of Ndc80C (WT or M1-M3 mutants) and 4 μM of CDK1-phosphorylated SkaC were combined in size exclusion chromatography (SEC) buffer (50 mM HEPES pH 8.0, 200 mM NaCl, 2.5 % glycerol, 1 mM TCEP) to a final volume of 65 μL. The samples were at least incubated for 1 h on ice or overnight at 10 ºC, and centrifuged before applying 50 μL of sample to the Superose 6 Increase 5/150 GL SEC column (Cytiva) pre-equilibrated with SEC Buffer. 80 μL fractions were collected, and the absorbance of the protein samples were monitored at 280 nm and 260 nm. Peak fractions were analyzed on 12 % Tricine-SDS-PAGE gel followed by Coomassie staining.

<u>Cell culture</u>

HeLa cells were cultured in DMEM (PAN-Biotech) supplemented with 10 % tetracycline-free FBS (PAN-Biotech), 2 mM L-Glutamine (PAN-Biotech), and 50 μg/mL Penicillin/Streptomycin (PAN-Biotech). Cells were grown at 37 °C with 5 % $CO_2$. Transfections and electroporations were carried out using DMEM without Penicillin/Streptomycin. Trypsinization was achieved using Trypsin/EDTA 0.05 %/0.02 % in DPBS (PAN-Biotech) for 5 min at 37 °C.

siRNA transfection

Depletion of endogenous Ndc80C was achieved using Lipofectamine RNAiMAX Transfection Reagent (Thermo Fisher Scientific) according to manufacturer's instructions and the following oligonucleotides: 60 nM siNdc80C (siNDC80 5′-GAGUAGAACUAGAAUGUGA-3′; siSPC24 5′-GGACACGACAGUCACAAUC-3′; siSPC25 5′-CUACAAGGAUUCCAUC AAA-3′, Sigma-Aldrich) for 48 h. Ndc80C depletion was obtained with a double transfection: all oligos were first added to the cells with reverse transfection. The treatment was repeated on adherent cells after 24 h (forward transfection).

Electroporation of recombinant proteins in human cells

For complementation assays a Neon Transfection System and Kit (Thermo Fisher Scientific) were used. HeLa cells depleted of the endogenous Ndc80C were trypsinized, washed in PBS and resuspended in electroporation buffer R (electroporation slurry 115 μL) (Thermo Fisher Scientific). Recombinant Ndc80 complexes (WT and mutants) were added to the slurry at a final concentration of 8 μM. HeLa cells were electroporated applying the following parameters: 2 pulses of 35 ms at 1000 V. After electroporation cells were resuspended in 13 mL warm PBS and centrifuged for 5 min at 500 x g. Next, cells were incubated for 5 min at 37 °C with 5 mL of trypsin/EDTA and subsequently washed twice with PBS (previously described in (*39*)). Subsequently, cells were resuspended in 1 mL DMEM, incubated for 3 min at 37 °C and seeded on poly-L-lysine (Sigma-Aldrich) coated coverslips in 12-well plates. Following 8 h of recovery, cells were treated with 9 μM RO3306 (Calbiochem) and incubated for 16 h. RO3306 was washed out three times with pre-warmed PBS and once with pre-warmed complete DMEM. Subsequently cells were treated with 10 μM MG132 (Calbiochem) for 1 h.

Cold treatment and immunofluorescence

Cold treatment was achieved by placing 12-well plates on ice and replacing the culture media with complete DMEM at 4 °C for 10 min. Untreated and cold-treated cells were permeabilized for 5 min before fixation with a solution of 0.5% Triton X-100 in PHEM buffer (PIPES, HEPES, MgSO4, EGTA) supplemented with 100 nM microcystin. Next, cells were fixed in 4 % paraformaldehyde (PFA) in PHEM buffer for 20 min and blocked with 5 % boiled goat serum (BGS) in PHEM buffer for 1 h. Coverslips were incubated for 2 h at room temperature with the following antibodies: CREST/anti-centromere antibodies (human, Antibodies Inc., 1:200), anti-CENP-C (guinea pig polyclonal, MBLPD030, MBL, 1:1000), anti-HEC1 (mouse, clone 9G3, Gene-Tex Inc., 1:1000), anti-SkaC (rabbit, generated in house, 1:5000), anti-α tubulin (mouse, clone DM1A, Sigma-Aldrich, 1:1000). Coverslips were washed three times for 5 min with a solution of 0.1 % Triton X-100 in PHEM buffer (PHEMT). Subsequently, cells were incubated for 1 h at room temperature with the following secondary antibodies: anti-human Alexa Fluor 647 (Jackson ImmunoResearch, 1:200), anti-mouse Rhodamine Red (Jackson ImmunoResearch, 1:200), anti-rabbit Alexa Fluor 488 (Jackson ImmunoResearch, 1:200). DNA was stained with 0.5 μg/mL DAPI (Serva) in concomitance with secondary antibodies. Coverslips were washed with PHEMT as previously described and rinsed with water before mounting with Mowiol (Calbiochem). Primary and secondary antibodies were diluted in 2.5 % BGS in PHEMT buffer.

Immunofluorescence imaging

Cells were imaged using a spinning-disk confocal device on the 3i Marianas system equipped with an Axio Observer Z1 microscope (Zeiss), a CSU-X1 confocal scanner unit (Yokogawa Electric

Corporation), 100 ×/1.4 NA Oil Objective (Zeiss), and Orca Flash 4.0 sCMOS Camera (Hamamatsu). Images were acquired as z sections of 0.27 µm using Slidebook Software 6 (Intelligent Imaging Innovations). Cells were imaged at room temperature. Images shown in figures (**Figure 3**F, **Figure S5**A, E-F) were converted into maximal intensity projections, exported, cropped, adjusted and converted into 16-bit TIFF files. Figures were arranged with Adobe Illustrator 2026. Quantifications of single kinetochore signals were performed using the software Fiji with an in-house macro as follows: CENP-C was used as reference on four-channels maximal intensity projections to generate two concentric spot-like circular ROIs (small 7 pixel, large 9 pixel of diameter) on single kinetochores. Each spot was analyzed individually and the final signal intensity per channel was obtained by subtracting the local background ("donut" shape around the smaller ROI) from the total signal intensity. Measurements were exported in Excel (Microsoft) and graphed with GraphPad Prism 9.0 (GraphPad Software).

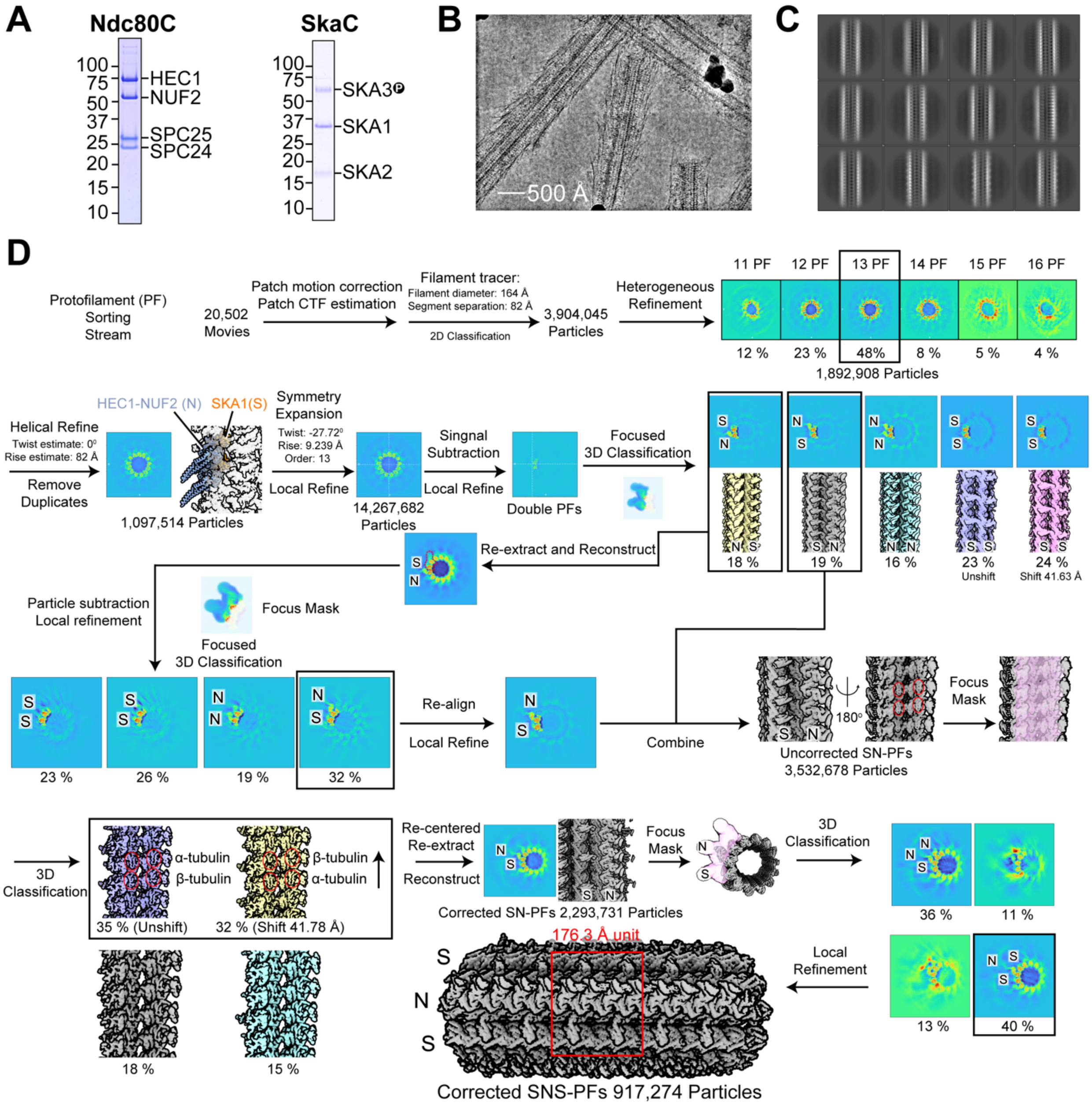


**Figure S1. Cryo-EM sample preparation and data processing.**
**(A)** SDS-PAGE analysis of purified Ndc80C-WT and pre-phosphorylated SkaC-WT used in this study.
**(B)** Representative cryo-EM micrograph of Ndc80C and SkaC bound to microtubules. Scale bar, 500 Å.
**(C)** Representative 2D class averages of selected 13-protofilament (PF) particles used for initial helical refinement in (D).
**(D)** Cryo-EM data-processing workflow illustrating the PF-sorting strategy used to identify homogeneously decorated SNS-PFs.

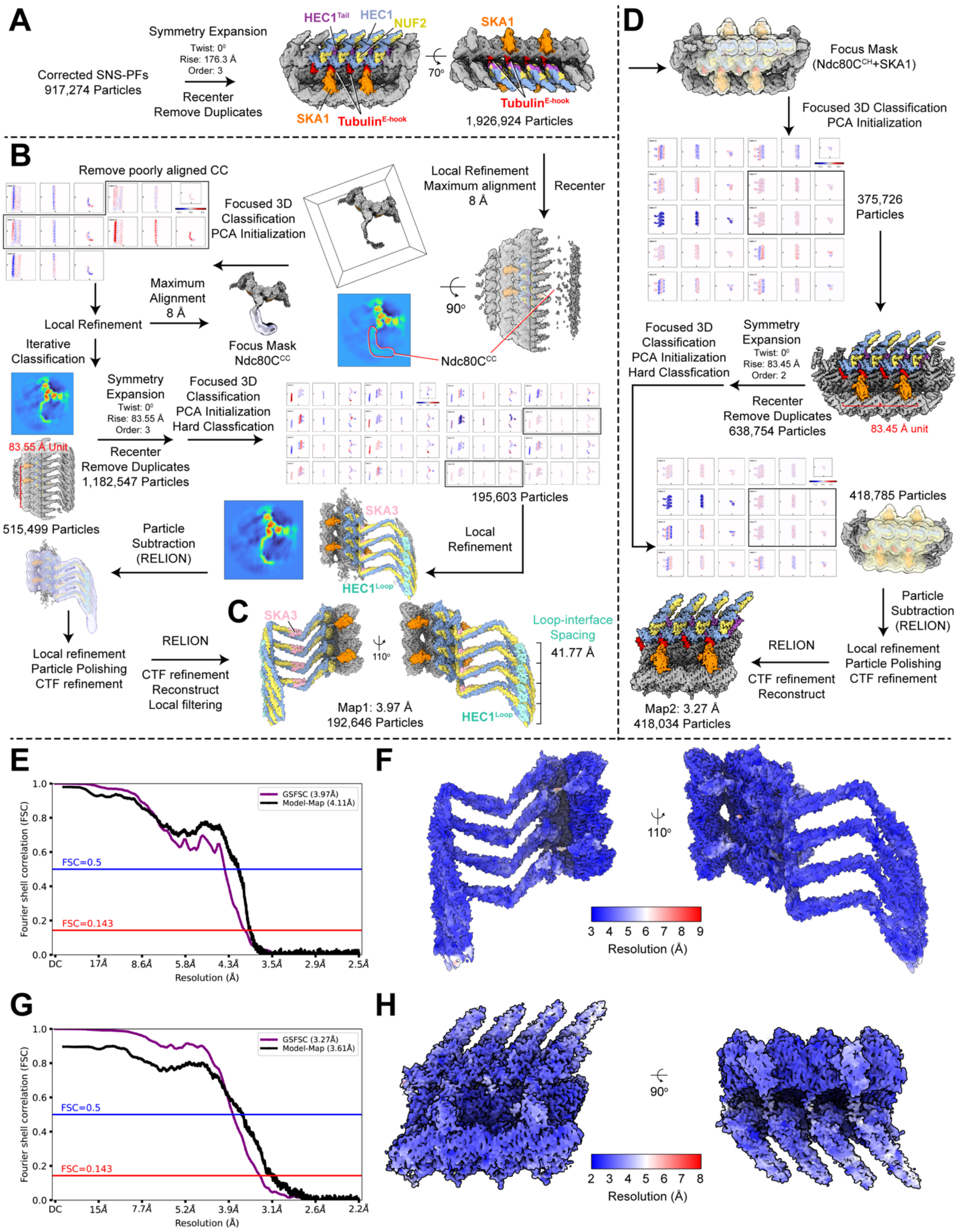


**Figure S2: Cryo-EM data-processing workflows for reconstructions of the SKA3, HEC1-kink, HEC1-loop interface and SKA1, HEC1-tail and microtubule interface.**

**(A)** Consensus reconstruction of selected SNS-PF particles from **Figure S1**D with a 176.3 Å repeating unit. Resolved Ndc80C and SkaC subunits are labeled and colored: HEC1 (blue), NUF2 (yellow), and SKA1 (orange). At lower contour levels, densities corresponding to the HEC1 tail (purple) and tubulin E-hooks (red) are visible.
**(B)** Cryo-EM data-processing workflow used to reconstruct the SKA3, HEC1-kink, HEC1-loop interface (map 1) by extensive 3D classification, signal subtraction, and local refinement approaches.
**(C)** Cryo-EM density map of the SKA3-HEC1 kink/loop interface (map 1). HEC1 (blue), NUF2 (yellow), SKA1 (orange), and SKA3 (pink) are shown. The HEC1 loop interface is highlighted in cyan and the ~41.77 Å spacing between adjacent HEC1 loops is indicated.
**(D)** Cryo-EM data-processing workflow used to reconstruct the SKA1, HEC1-tail, microtubule interface (map 2) using extensive 3D classification, signal subtraction, and local refinement approaches.
**(E)** Fourier shell correlation (FSC) curves for the deposited map of the SKA3, HEC1-kink, HEC1-loop interface (map 1) and the corresponding model-to-map FSC.
**(F)** Local resolution estimation for map 1 to reflect the overall quality of the final reconstructed map.
**(G)** FSC curves for the deposited map of the SKA1, HEC1-tail and microtubule interface (map2) and the model-to-map FSC curve.
**(H)** Local resolution estimation of map2 to reflect the overall quality of the final reconstructed map.

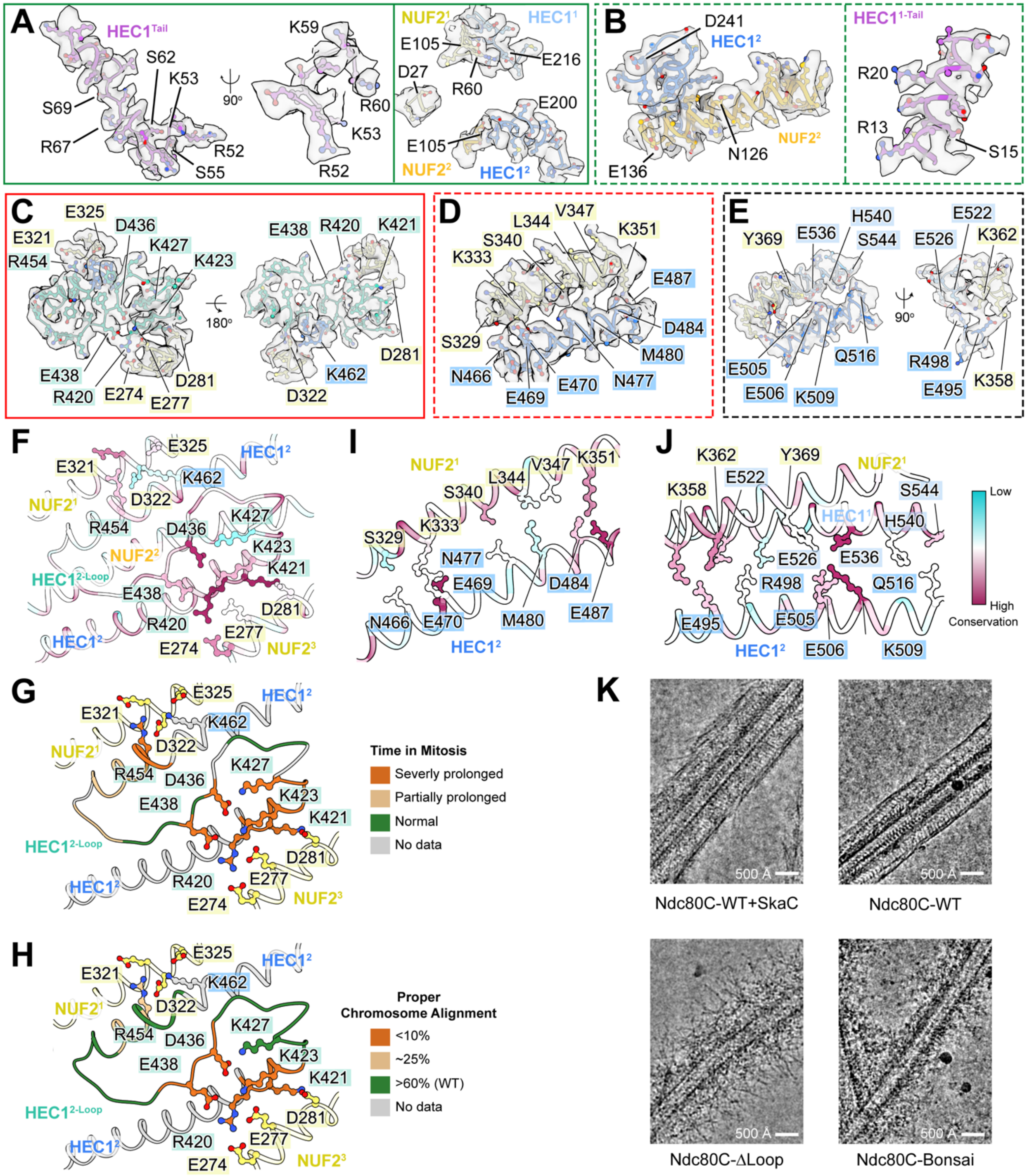


**Figure S3: Structural analysis of the longitudinal oligomerizations of Ndc80C mediated by the HEC1-tail and loop interfaces.**

**(A-B)** Cryo-EM density showing the HEC1 tail bridging adjacent Ndc80C CH domains, corresponding to **Figure 2**C (A) and **Figure 2**D (B). For better visualization, viewing angles match those in **Figure 2**C and 2D, while HEC1 tail density and the corresponding Ndc80C CH domains are shown separately.

**(C-E)** Cryo-EM density of the HEC1 loop interface (C, corresponding to **Figure 2**F) and contacts formed by the HEC1 and NUF2 coiled-coils (D and E, corresponding to **Figure 2**G and 2H). Left panels in (C) and (E) show views matching **Figure 2**F and 2G, respectively; right panels show rotated views. The viewing angle in (D) matches **Figure 2**G.
**(F)** Sequence conservation analysis of the HEC1 loop interface (F, corresponding to **Figure 2**F) Residues are colored from variable (cyan) to conserved (magenta) based on sequence alignments.
**(G-H)** Summary of functional analyses of residues at the HEC1 loop interface, viewed from the same angle as **Figure 2**F. **(G)** Data from (*29*), in which alanine substitutions caused severe, partial, or no mitotic delay (orange, brown, and green, respectively). **(H)** Data from (*19*), in which alanine substitutions resulted in <10 %, ~25 %, or >60 % proper chromosome alignment at metaphase (orange, brown, and green, respectively). Residues lacking functional data are shown in gray.
(**I**-**J**) Sequence conservation analysis of the coiled-coil contacts between HEC1 and NUF2 (I and J, corresponding to **Figure 2**G and 2H), viewed from the same angles as **Figure 2**G-2H. Residues are colored from variable (cyan) to conserved (magenta) based on sequence alignments.
**(K)** Representative cryo-EM micrographs of microtubules decorated with Ndc80C-WT and SkaC-WT, Ndc80C-WT alone, Ndc80C-ΔLoop, or Ndc80C$^{Bonsai}$ (lacking the HEC1 loop and most coiled-coil regions). Scale bar, 500 Å.

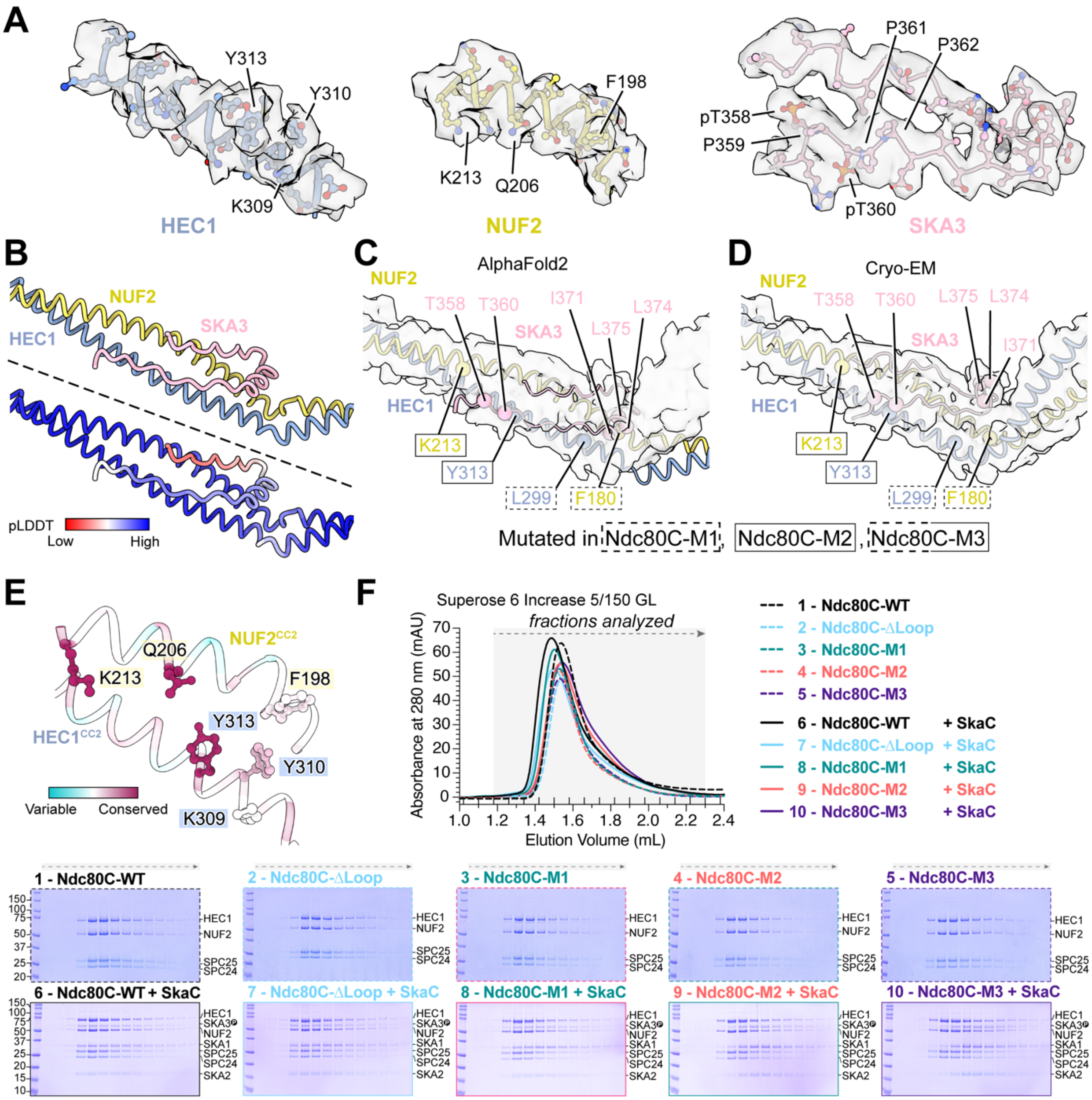


**Figure S4. Characterization of SKA3-binding interface of Ndc80C.**

**(A)** Cryo-EM density of HEC1, NUF2, and SKA3 at the Ndc80C-SkaC interface, corresponding to **Figure 3**C. The viewing angle is held constant, with individual components shown separately for clarity.

**(B)** AlphaFold2 prediction of the Ndc80C-SKA3 interaction, with corresponding per-residue confidence (pLDDT) scores.

**(C-D)** Comparison of the Ndc80C-SKA3 binding interface from AlphaFold2 (C) and cryo-EM (D), with residues mutated in Ndc80C highlighted (M1: HEC1 L299A, NUF2 F180A; M2: HEC1 Y313A, NUF2 K213Q; M3: combination of M1 and M2, respectively).

**(E)** Sequence conservation analysis of the Ndc80C-SkaC binding interface, viewed from the same angle as **Figure 3**C. Residues are colored from variable (cyan) to conserved (magenta) based on sequence alignments.

**(F)** Analytical size-exclusion chromatography of pre-phosphorylated SkaC interaction with Ndc80C variants (WT, ΔLoop, M1, M2, and M3) using a Superose 6 Increase 5/150 GL column. Retention volumes and corresponding SDS-PAGE analyses are shown. Sample 1, 6 and 9 correspond to the same experiment as shown in **Figure 3**D.

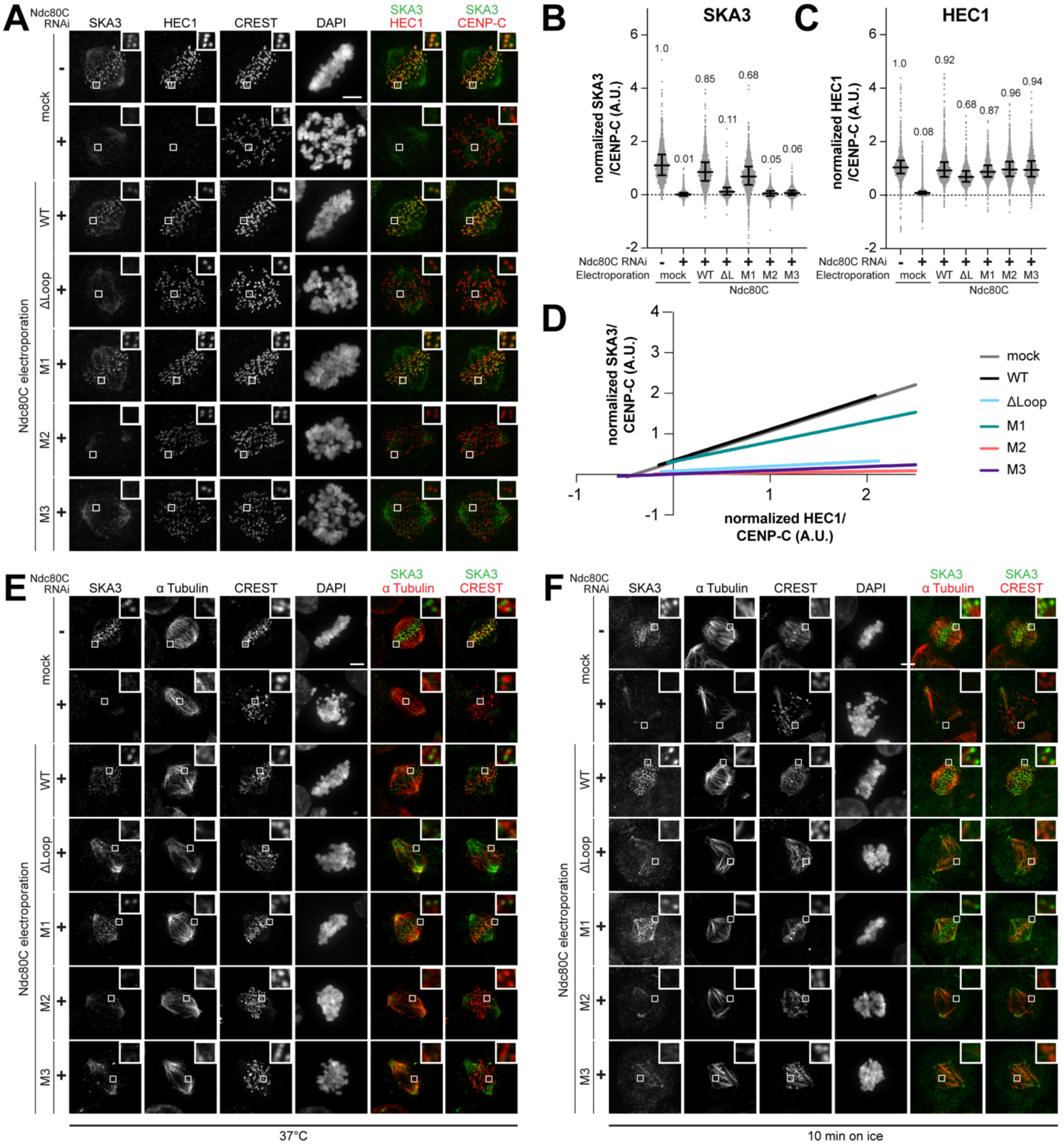


**Figure S5: In vivo validation of Ndc80C mutants.**

**(A)** Immunofluorescence of representative HeLa cells untreated or treated with Ndc80C RNAi and electroporated as indicated (mock, Ndc80C-WT, -ΔLoop, -M1, M2 or -M3). CENP-C is a kinetochore marker. Scale bar: 5 µm.

**(B-C)** Quantification of kinetochore levels of endogenous SKA3 (B) and endogenous or electroporated HEC1 (C) normalized to endogenous levels (no RNAi). Number of kinetochores analyzed from three independent experiments: n = 2669 for control, n = 2773 for RNAi, n = 2508 for WT, n = 1563 for ΔLoop, n = 3037 for M1, n = 2506 for M2 and n = 2566 for M3. Black bars

represent median, indicated above the plots, and interquartile range of normalized single kinetochore intensities.
**(D)** Linear regression of data points for each kinetochore. Normalized SKA3 and HEC1 intensities are plotted on y- and x-axis, respectively.
**(E-F)** Immunofluorescence of representative HeLa cells untreated or treated with Ndc80C RNAi, electroporated as indicated (mock, Ndc80C-WT, -ΔLoop, -M1, M2 or -M3) and either directly fixed or cooled for 10 min and then fixed. Scale bar: 5 µm.

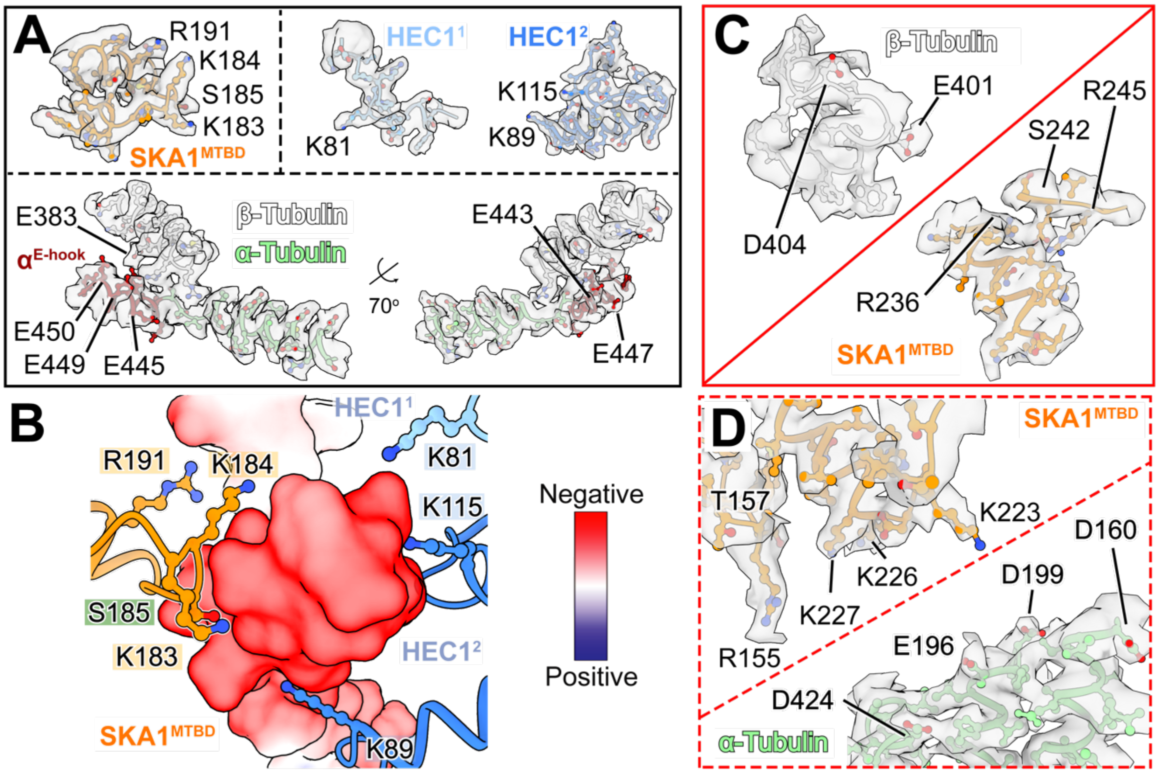


**Figure S6: Structural analysis of microtubule binding of Ndc80 and Ska complexes.**
**(A)** Cryo-EM density for binding interfaces of SKA1-MTBD with oligomerized Ndc80C by the E-hook of α-tubulin from PF2 (A, corresponding to **Figure 4**C). For better visualization, viewing angles for SKA1$^{MTBD}$, HEC1$^{1}$ and HEC1$^{2}$ are consistent to **Figure 4**C (top panel), while those for the tubulin are rotated (bottom panel).
(**B**) Representation of surface-charged distribution of α-tubulin E-hook and β-tubulin interfaced between SKA1$^{MTBD}$, HEC1$^{1}$, and HEC1$^{2}$ chains as shown in **Figure 4**C. Putative Aurora B phosphorylation site found in SKA1$^{MTBD}$ (S185) can be identified in closed proximity to binding interface.
**(C-D)** Cryo-EM density for binding interfaces of SKA1-MTBD with β-tubulin (C, corresponding to **Figure 4**D) or α-tubulin (D, corresponding to **Figure 4**E) from PF1, respectively. For better visualization, viewing angles are consistent compared to **Figure 4**D (C) and 4E (D) with different proteins shown separately.

**Table S1. Cryo-EM data collection, processing, and refinement statistics.**

| | SKA3, HEC1-kink and Hec1-loop interfaces[Map1] | SKA1, HEC1 tail and microtubule interface[Map2] |
|---|---|---|
| **EMDB ID** | EMD-XXXXX | EMD-XXXXX |
| **PDB ID** | XXXXX | XXXXX |
| **Data collection** | | |
| Microscope | Titan Krios G2 | |
| Detector | K3 summit | |
| Voltage (kV) | 300 | |
| Pixel size (Å) | 0.44 | |
| Total electron exposure ($e^-/Å^2$) | 58 | |
| Defocus range (μm) | -1.5 to -2.3 | |
| Micrographs collected | 20,502 | |
| **Reconstruction** | | |
| Final particle images | 192,646 | 418,034 |
| Pixel size (Å) | 1.232 | 1.1 |
| Box size (pixels) | 400 | 320 |
| Resolution (Å) (FSC = 0.143) | 3.97 | 3.27 |
| Map Sharpening B-factor ($Å^2$) | -43.57 | -67.20 |
| **Model composition** | | |
| Non-hydrogen atoms | 72,724 | 59,957 |
| Protein residues | 9,017 | 7,467 |
| Ligands | 18 | 18 |
| Metals | 12 | 12 |
| **Refinement** | | |
| Resolution (Å) (FSC = 0.5) | 4.11 | 3.61 |
| Model-to-map CC (mask) | 0.82 | 0.73 |
| Model-to-map CC (volume) | 0.82 | 0.73 |
| R.m.s deviations | | |
| Bond length (Å) | 0.004 | 0.006 |
| Bond angles (°) | 1.038 | 1.062 |
| **Validation** | | |
| MolProbity score | 1.21 | 1.10 |
| Clash score | 3.23 | 2.86 |
| Ramachandran plot | | |
| Outliers (%) | 0.0 | 0.0 |
| Allowed (%) | 2.48 | 2.08 |
| Favored (%) | 97.52 | 97.92 |
| Rotamer outliers (%) | 0.40 | 0.85 |
| C-beta deviations (%) | 0 | 0 |
